# Enhanced spatially resolved transcriptomics analysis by matching between expression profiles and spatial topology

**DOI:** 10.1101/2024.08.16.608230

**Authors:** Yuxuan Pang, Chunxuan Wang, Yao-zhong Zhang, Zhuo Wang, Seiya Imoto, Tzong-Yi Lee

## Abstract

Spatially resolved transcriptomics (SRT) quantifies gene expression covering contextual heterogeneity for the tissue section. Exploratory data analysis using SRT data sheds light on diverse biomedical research fields. We propose STForte, a pairwise graph autoencoding-based approach for SRT data analysis, which is capable of matching the information between expression profiles and spatial topology in the latent space. STForte benefits from the designed framework to provide encodings with justifiable spatial correlations for the down-stream analysis of both homogeneous and heterogeneous SRT data. Moreover, STForte can unravel the biological patterns of unobserved locations or recover deficient measurements to enable spatial enhancement. Latent encodings generated by STForte can be used to perform spatial region identification or other downstream tasks to gain biological insights and elucidate biological processes. In this work, we presented various analyses to show that STForte is scalable for analyzing SRT data under different scenarios with considerable performance.

## 1 Introduction

To comprehend the functioning of life, it is crucial to observe the behavior of cells or groups of cells in multicellular organisms, even though they share a common genome, as their morphology and gene expression patterns show unique and changing differences [1]. One approach to achieving this is through the use of spatially resolved transcriptomics (SRT) [2], which aims to quantify the mRNA expression of a number of genes within the spatial organization of tissues to illustrate both the positional and contextual information of the transcripts [3]. It provides an articulate view for specifying diversity among different spatial regions, thereby understanding crucial biological processes.

Recent advances in SRT have enabled researchers to test hypotheses and uncover insights through observation and exploratory data analysis (EDA)[4]. To this end, various computational methods have been proposed to facilitate the progress of EDA using SRT data. Specifically, latent encoding extraction and spatial region identification are essential processes in SRT analysis that provide condensed information or articulate demonstrations related to tissue domains for subsequent downstream analyses. These methods can be based on statistical frameworks[5–7] or follow graph neural network architectures [8–12]. Most existing methods consider introducing spatial correlations between neighboring spots to induce spatial homogeneity, thereby specifying coherent patterns for spatial regions [13]. Ascertaining spatial homogeneity in SRT analysis benefits related tasks in dissecting consecutive spatial domains, such as identifying anatomical regions for embryogenesis or nervous system discovery. However, excessive consideration of spatial correlation may cause overemphasis on homogeneous and smooth information, resulting in the absence of heterogeneous patterns. This could be defective in analyzing complex tissues with high heterogeneity, such as tumor micro-environments [14–16].

SRT technologies can generally be categorized into two types [17]: imaging-based and next-generation sequencing (NGS)-based technologies. Imaging-based approaches [18–22] use probe-based identification to chart the precise locations of a selected set of transcripts to provide high-resolution information on cellular patterns and greater depth for capturing transcripts. However, they are constrained by the need for pre-selected genes, which results in a limited and biased field of view. In contrast, NGS-based technologies [23, 24] perform unbiased mRNA capture by barcoding transcripts from predetermined positions (spots) to obtain gene expression profiles, which are more accessible with standardized protocols and are unbiased in measuring genes. However, these methods are still restricted by their relatively low sensitivity and resolution. Specifically, although there are recently flourished NGS-based approaches with increasingly higher spatial resolution in transcript measurements [25–28], some of the trending approaches [29–31] have limited resolution and spatial coverage in capturing the transcripts of intact tissue, resulting in inconsistent spatial patterns of transcript measurements. Therefore, computational methods have been proposed to facilitate SRT analysis through in silico enhancement of spatial resolution. Resolution enhancement can be performed by combining histological images [32, 33] to predict pixel-level expression profiles for tissue sections or by considering breaking the measured spots into sub-spot resolutions using modified probabilistic-based approaches [6, 7]. Besides, another approach considers leveraging methods from computer vision, utilizing deep neural networks to directly predict expression values for predefined unobserved locations [34]. Nevertheless, the existing methods either depend on histological images and large computational resources or are deficient in flexibility and the consideration of structural contexts for spatial imputation.

Here, we propose STForte, a graph-learning-based autoencoding approach, to address the aforementioned challenges in SRT data analysis. STForte is based on a modified pairwise graph autoencoder-based architecture specified by deep neural networks, which is crafted to simultaneously handle the gene expression profiles and spatial topology information of SRT data by manipulating their correspondence within the latent space. On this basis, STForte highlights the following aspects. First, modeling the correspondence enables the framework to adaptively capture the impartial patterns between the expression attributes and spatial correlation, which facilitates the capability in characterizing both the homogeneity and heterogeneity in SRT data analysis. Second, the spatial topology is encoded to unify the spatial correlation, which can be used to impute the biological patterns of locations with unmeasured or deficient transcripts. With the designed procedures, the encoded spatial topology can be used to perform spatial enhancement to provide analyses with more consecutive content and fine-grained resolution. Third, STForte is scalable to accommodate various analytical scenarios for SRT data. The generated latent encodings can be employed to conduct spatial region identification or even more advanced downstream tasks in combination with other methods, including but not limited to spatial trajectory inference [35] and cell-cell interactions [36]. Extensive experiments were conducted, demonstrating that STForte is applicable to various SRT data obtained from imaging-based (e.g., 10x Xenium) and NGS-based (e.g., 10x Visium, Stereoseq) platforms, and can be adapted for single-slice and multi-slice analysis. We believe that STForte can reinforce SRT data analysis in a more comprehensive and fine-grained view to discover contextual biological insights through spatially resolved transcriptomics.

## 2 Results

### 2.1 Overview of STForte method

The schematic overview of STForte is illustrated in Fig. 1, with technical details described in “Method” and Supplementary Notes. Briefly, SRT data are processed to extract their topological relations and transcription profiles (expression attributes). For some NGS-based SRT data protocols with latticed technical specs, such as 10x Visium, STForte can automatically generate new padding spots from the original collection of spots based on their coordinates and predefined distance. Users can also manually mask spots for mismeasured or unmeasured locations to infer their biological patterns. A topological graph is subsequently established using spatial k-nearest neighbors (KNN) or their straightforward closeness based on a predefined distance. For the observed expression attributes, STForte can directly accept a preprocessed count matrix or, alternatively, use results obtained through unstructured dimensionality reduction methods such as PCA or scVI [37].

**Fig. 1.**
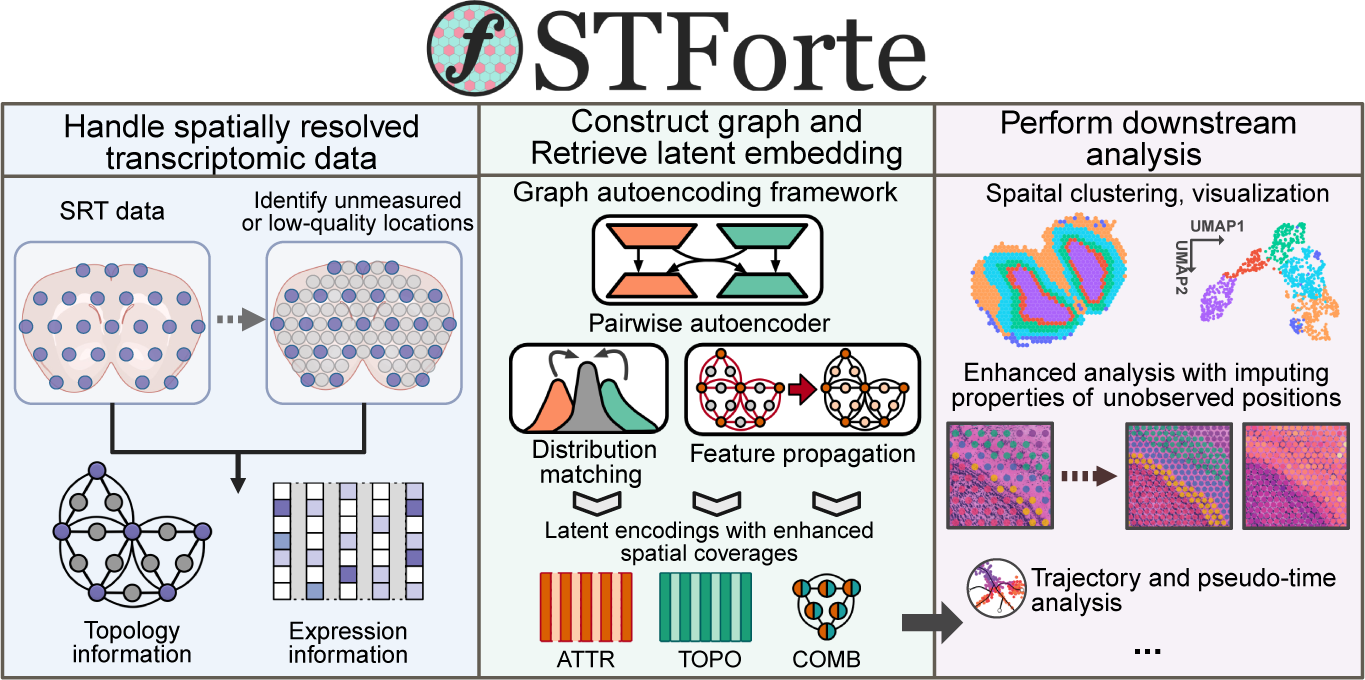
Schematic overview of our proposed approach. STForte first processes SRT data to extract spatial topology and expression attributes. It can also identify unmeasured or low-quality locations through automatic or user-defined procedures, which can be included in subsequent analyses. STForte uses a pairwise graph autoencoder-based structure to learn low-dimensional latent encodings based on expression attributes (ATTR) and topological graph (TOPO). Distribution matching and feature propagation are two modules that facilitate STForte in inferring encodings for unmeasured or low-quality locations. These latent encodings, along with their combination (COMB), can be used for various analyses such as identifying spatial domains, UMAP visualization, and spatial trajectory analysis. Moreover, inferred encodings for unmeasured or mismeasured locations can lead to improved spatial consistency and resolution in analysis.

STForte adopts a pairwise graph autoencoder (GAE)-based framework (Extended Data Fig. 1) with adversarial distribution matching to jointly learn the latent spaces for both the expression attributes and spatial topology information of SRT data. The pairwise GAE framework also enables STForte to infer the biological patterns of locations with inaccessible or mismeasured transcripts through latent encoding using an approach of learning on the attribute-missing graph. In addition, a feature propagation technique is adopted to facilitate the inference of inaccessible locations for expression attributes. Subsequently, the attributes (ATTR), topology (TOPO) encodings, and their combined encodings by concatenation (COMB) can serve diverse downstream analyses, such as visualization and spatial region identification through clustering algorithms (e.g., mclust, Leiden, or Louvain) or trajectory analysis through PAGA [35]. Moreover, inferred latent encodings can be used for propagating spatial region annotation or predicting unseen expression values for padding spots or mismeasured locations, which enables analysis and discovery under better resolution or in extra biological contexts.

### 2.2 STForte facilitates analysis with refined resolution and consistent patterns on 10x Visium mouse olfactory bulb data

We first applied STForte to a mouse olfactory bulb dataset from the 10x Visium platform [38]. The hematoxylin and eosin(H&E)-stained image of the dataset in Fig. 2a shows the laminar organization of the coronal mouse olfactory bulb, which is composed of granule cell layer (GCL), glomerular layer (GL), external plexiform layer (EPL), mitral cell layer (MCL), and olfactory nerve layer (ONL). As the 10x Visium platform has locations of void measurement among the lattice arrangement of spots, the padding strategy was performed in the preprocessing step by inserting new unobserved spots for subsequent inference (Extended Data Fig. 2a) and used STForte to extract three latent encodings (ATTR, TOPO, and COMB) for the data. Gaussian mixture model (GMM)-based clustering was conducted using mclust to identify the spatial regions. As presented in Fig. 2b, The results show that the laminar structure is well recognized by the clustering results of COMB encoding, while its UMAP visualization has a consistent transition from the outer ONL to the inner GCL. Opposed to COMB, despite ATTR encoding having a slightly higher silhouette coefficient and Calinski-Harabasz (C-H) index, it failed to provide consistent organization of the laminar structure, while TOPO encodings have consistent organization but fall short in distinguishing between ONL and GL (Extended Data Fig. 2b, c). Subsequently, we propagated the spatial region identification result from the observed to padding spots using three different encodings. Here, TOPO encoding provides a more refined organization and distinct boundaries for laminar structure (Fig. 2c). Specifically, the MCL layer in the original view (Fig. 2b) is relatively inconsistent owing to the limited spatial resolution of SRT technology, whereas STForte helps to recover more consistency and fine-grained organization through the padding view. In contrast to TOPO, both ATTR and COMB encodings are relatively deficient in giving coherent structures under the padding scenario (Extended Data Fig. 2d).

**Fig. 2.**
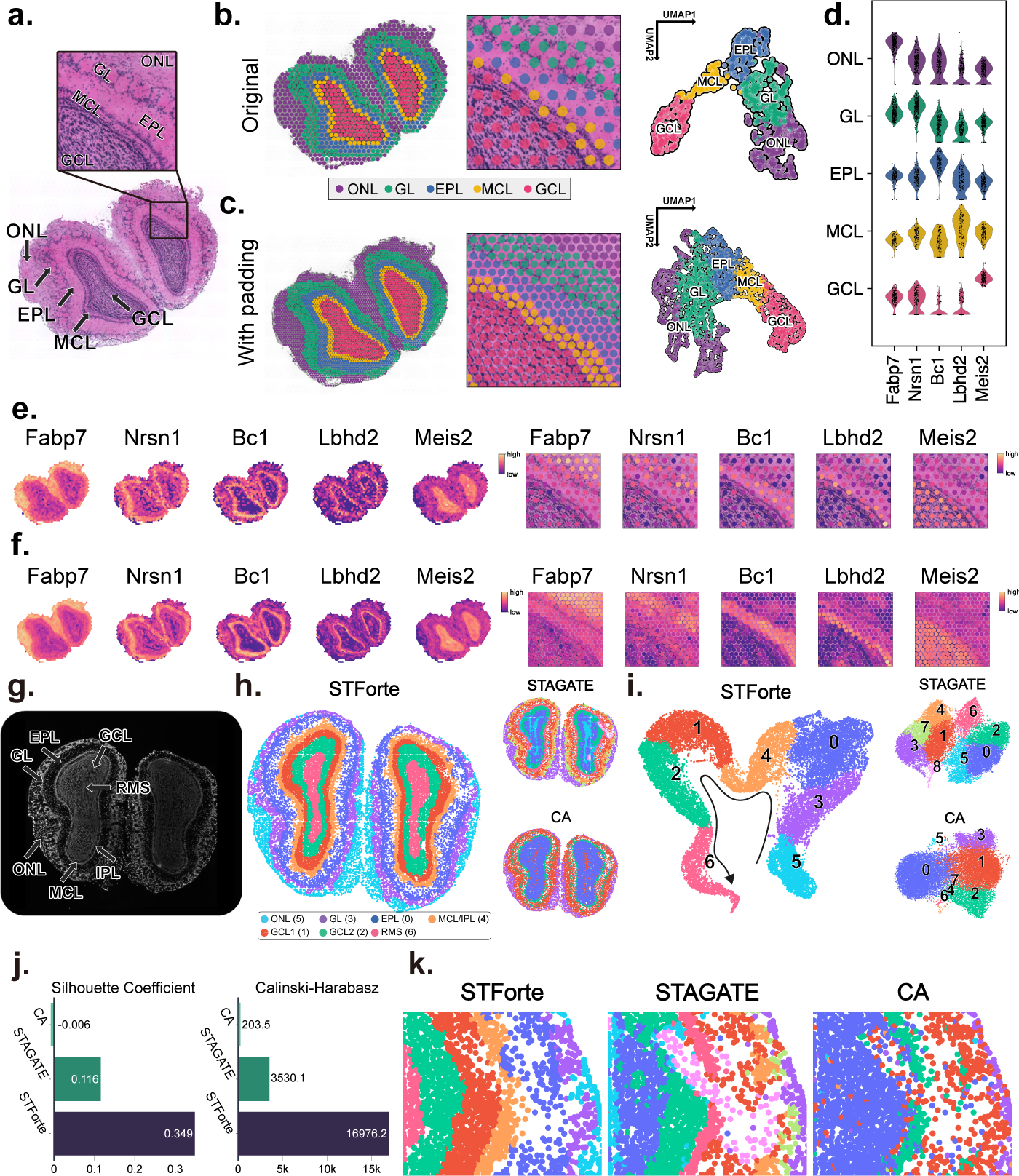
**a,** H&E-stained image and annotation of the 10x Visium mouse olfactory bulb data including a zoomed-in region. **b,** Spatial regions and UMAP visualization obtained from STForte (COMB encoding). **c,** Spatial regions and UMAP visualization under padding views obtained from STForte (TOPO encoding). **d,** Violin plots of gene expression levels of the selected 5 layer-specific marker genes. **e,** Visualization of the spatial expression levels of five layer-specific genes on the entire dataset (left) and the zoomed-in region (right). **f,** Spatial expression levels after STForte’s padding on the entire dataset (left) and the zoomed-in region (right). **g,** DAPI-stained image with manual annotation of the Stereo-seq mouse olfactory bulb data. **h,** Spatial regions identified by different dimensional reduction methods, including STForte, STAGATE and CA. **i,** UMAP visualizations obtained by different methods. **j,** Comparison of clustering metrics for different methods. **k,** Zoomed-in region of interest showing the results of low-quality spots processed by different methods.

Next, we examined some layer-specific marker genes according to the spatial region annotations recognized by STForte. The results showed that they were expressed at relatively high levels within their corresponding layers (Fig. 2d, e, Supplementary Fig. S1 and additional Wilcoxon rank-sum test results in Supplementary Table S1), which is concordant with previous reports or validations. For instance, *Fabp7* is highly expressed in the identified ONL, where it plays a crucial role in the development, maintenance, and function of this specialized sensory tissue, which is responsible for detecting odors and transmitting olfactory information to the brain [39]. *Nrsn1* is highly expressed in the GL, where it contributes to olfactory signal processing and transmission via its involvement in synaptic vesicle regulation, axonal targeting, and neuroplasticity [40]. *Bc1*, a non-coding RNA that plays a crucial role in facilitating the refining and processing of olfactory signals, is abundant in the EPL [41]. The *Lbhd2* gene, which has been employed to achieve more precise genetic targeting of mitral cells, also exhibited high expression levels in the identified MCL [42]. As a typical genetic marker for interneurons in the olfactory bulb, *Meis2* [43] exhibits a high expression pattern in the GCL derived from the STForte results. Furthermore, the application of STForte encodings can facilitate the extrapolation of gene expression levels by propagating to unmeasured locations. The resulting padding visualizations (Fig. 2f) markedly enhance the understanding of spatial organization, offering a more consistent and detailed perspective of spatial expression patterns. Additionally, we evaluated the spatial region identification of the dataset using STAGATE, a standard dimensionality reduction method designed for SRT data, as well as Correspondence Analysis (CA) [44], which is commonly used for reducing dimensions in regular gene expression data. We then compared these approaches with our method (Extended Data Fig. 2e). STForte achieved the highest silhouette coefficient, while STAGATE recorded the highest C-H index. The spatial region and UMAP visualizations of STForte, STAGATE, and CA are illustrated in Extended Data Fig. 2f, g. STAGATE identified a consistent laminar structure but could not distinguish between the EPL and MCL layers. The result of CA was slightly similar to the results of the ATTR encodings from STForte, which failed to provide consistent organization of the laminar structure.

### 2.3 Analyzing spatial cell-resolution mouse olfactory bulb data from Stereo-seq using STForte

Next, we examined the capability of STForte on analyzing the SRT data at cellular resolution. We applied STForte to the mouse olfactory bulb dataset obtained from the Stereo-seq platform, which utilizes cell segmentation and aggregated UMI counts to acquire cell-binned SRT data [27]. Fig. 2g shows the DAPI-stained image with manual annotation of the laminar structure of the olfactory bulb, which is composed of GL, EPL, MCL, IPL (internal plexiform layer), GCL, and RMS (rostral migratory stream). In this dataset, certain spots exhibit poor expression measurement quality (Extended Data Fig. 3a). STForte’s padding strategy allows for the retention of low-quality spots by masking and inferring their biological patterns based on the spatial context of neighboring cells. We conducted spatial region identification using COMB encoding and compared the results with those from STAGATE and CA (Fig. 2h). STForte’s results revealed the most consistent spatial regions, accurately distinguishing the laminar structure of the mouse olfactory bulb. We further explored the expression of layer-specific genes [39, 41–43] within these regions (Extended Data Fig. 3b, c, and additional Wilcoxon rank-sum test results in Supplementary Table S2), reinforcing the validity of the STForte clustering outcomes. For example, in the GCL, *Pcp4* and *Gad1* exhibit relatively higher expression throughout the outer part of the GCL (cluster 4), which is generally in concordance with the findings mentioned in [45, 46]. Additionally, the RMS region identified by STForte also shows high expression of its corresponding dominant gene, *Mbp*[47]. In contrast, although STAGATE provided relatively accurate annotations, the consistency between regions is comparatively lower. As for CA, which does not consider spatial information, the overall regional distribution appears to be more scattered. Although it also separately identified ONL, MCL/IPL, and GCL, it confused GL, EPL, and RMS together. In the UMAP visualizations of the three methods (Fig. 2i), STForte presented a continuous transition from ONL to RMS across different regions, which is not observed in the UMAP visualizations of STAGATE and CA. We compared the clustering performance metrics of the latent space across different methods (Fig. 2j), in which STForte performed better in terms of both the silhouette coefficient and the C-H index. Furthermore, STForte accurately recovered the biological patterns of masked spots, consistently assigning them to their neighboring spatial regions (Fig.. 2k). In contrast, the other two methods failed to make additional inferences for these low-quality spots based on spatial context, leading to either their separate differentiation or confusion among different annotations.

### 2.4 Exploring tumor micro-environment of prostate adenocarcinoma SRT data with STForte

Leveraging SRT data for cancer analysis presents significant challenges. The invasive growth of tumors, which penetrates their original location and infiltrates the surrounding tissues, results in a mixture between the tumor and the normal area. Moreover, tumors are often accompanied by inflammatory responses and immune cell infiltration. These inflammatory regions do not always perfectly align with tumor cell areas, adding further complexity to the spatial distribution of the tumor micro-environment (TME) [14–16].

We analyzed the prostate adenocarcinoma data from the 10x Visium platform using STForte. Fig. 3a shows the H&E-stained images and pathological annotations. We employed STForte for spatial encodings and performed clustering using the Leiden method, varying its resolution to compare the silhouette coefficients (SC) (Extended Data Fig. 4a). ATTR encoding yielded higher SC and was more coherent with the pathological annotations (Extended Data Fig. 4b, c). We selected ATTR as the primary encoding, and the spatial regions are shown in Fig. 3b-left. C1, C2, and C6 corresponded to invasive carcinoma locations, whereas the other clusters characterized non-tumor regions. In addition, padding by TOPO provided spatially consistent results and fine-grained information (Fig. 3b-right, Extended Data Fig. 4d, e). We also compared the STForte (ATTR) results along with STAGATE and CA. Fig. 3c shows the SC and numbers of clusters at different resolutions of the Leiden method. STAGATE and CA had higher coefficients at lower resolutions, but lower SC when STForte reached its maximum or at the same number of clusters. Fig. 3d and Extended Data Fig. 4f present the spatial regions and UMAP visualizations of STAGATE and CA, respectively. At its highest silhouette coefficient, CA could not distinguish the entire invasive carcinoma area from the tissue structures. With the same number of clusters as STForte (#Clust = 11), CA was able to reflect the TME composition but had a lower silhouette coefficient. STAGATE, on the other hand, failed to accurately describe the TME or distinguish tumors from other tissues. A similar phenomenon occurred in STForte’s TOPO encoding, indicating that excessive consideration of spatial adjacency may lead to oversmoothing. However, STForte provides different encodings for a trade-off between expression and spatial information, thus alleviating this problem to a certain extent.

**Fig. 3.**
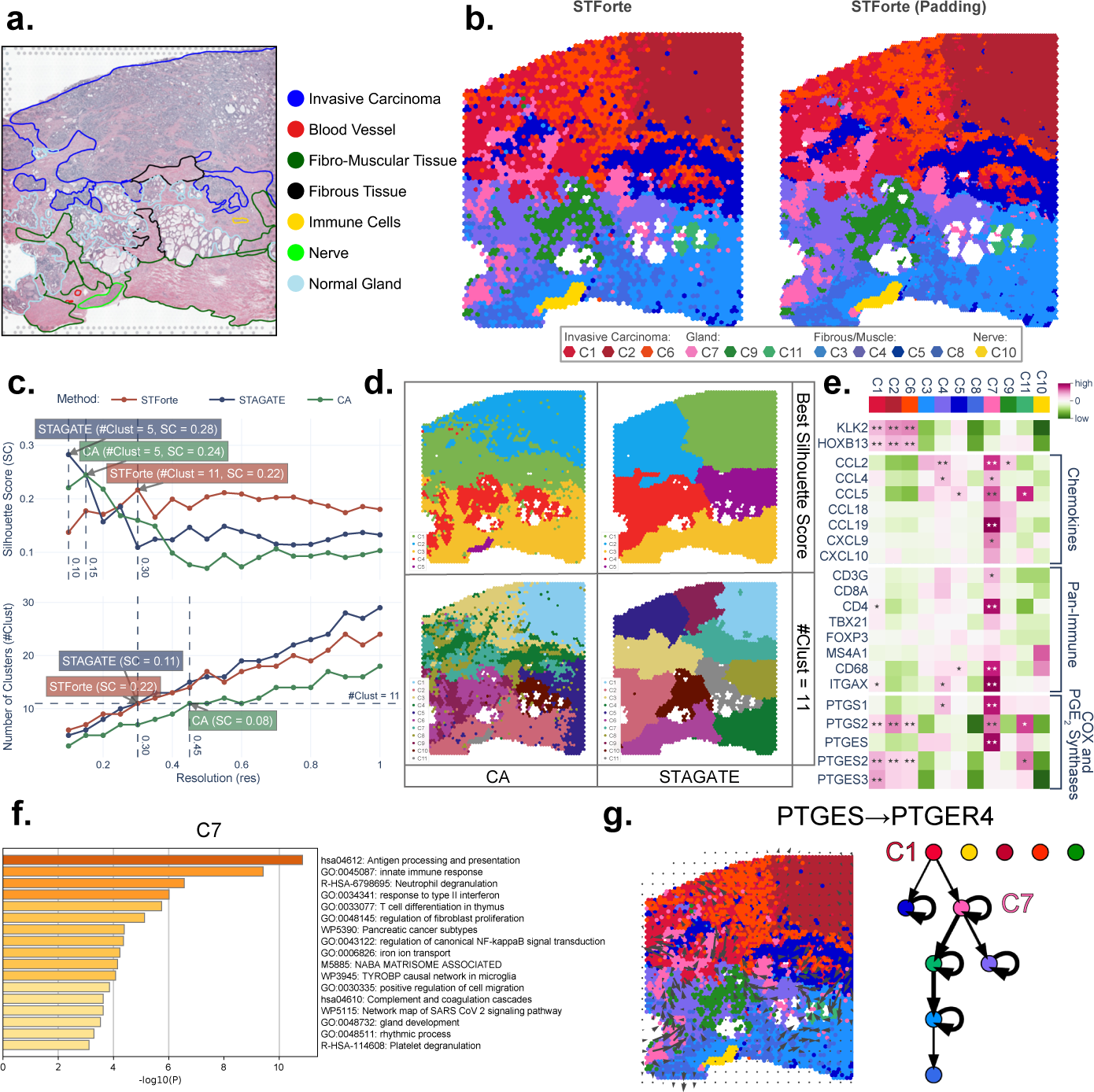
**a,** H&E-stained image with pathological annotations of the 10x Visium prostate adenocarcinoma. **b,** Spatial region identification using STForte ATTR encoding (left) and results from the padding scenario with STForte TOPO encoding (right). **c,** Silhouette coefficients (SC) of STForte, CA and STAGATE under the Leiden method at various resolutions. **d,** Spatial visualization shows the Leiden results from STAGATE and CA at their respective best silhouette coefficients or when #Clust=11. **e,** Heatmap depicting the mean expression levels of genes associated with cancer, immune response, inflammation, and PGE_2_ synthesis across distinct spatial regions. The statistical significance of higher relative expression within each region was assessed using one-sided Wilcoxon rank-sum tests. Stars denote significance levels: ★: p*<*0.05; ★★: p*<*0.001. **f,** Significant biological pathways associated with region C7 based on the differentially expressed genes. **g,** Spatial interaction between *PTGES* and *PTGER4* genes based on STForte-identified regions and COMMOT analysis.

Furthermore, we investigated the expression of canceror immune-related genes across different spatial regions identified by STForte (Fig. 3e, Supplementary Fig. S2). *KLK2* and *HOXB13* have been reported to be strongly associated with the proliferation and metastasis of prostate cancer cells [48–50], and their expression levels were elevated in the invasive carcinoma regions. Chemokines, such as *CCL2*, *CCL5*, and *CCL19*, which play crucial roles in immune cell recruitment and the inflammatory response [51, 52], were found to be relatively enriched in C7. *CD3G* is a marker of T cell differentiation and maturation [53, 54], whereas *CD4* is an important molecule expressed on the surface of helper T cells [55], and their corresponding genes were both observed to have significantly higher expression in C7. Additionally, the expression levels of *CD68* and *ITGAX* have relatively higher expression levels in C7. *CD68* is a glycoprotein mainly expressed on the surface of monocytes, macrophages, and other myeloid cell lineages [56], whereas *ITGAX* encodes the CD11c, which is a marker of dendritic cells [57]. To this end, we speculated that C7, identified by STForte, is a potential Tertiary Lymphoid Structure (TLS) [58, 59] that exhibits intense cancer-related immune inflammatory responses. We also investigated representative biological pathways for each spatial region based on the calculation of differential expressed genes (Supplementary Fig. S3). The significant pathways in regions related to invasive carcinoma mentioned the relevant effects of the androgen receptor and prostate cancer or *KLK2* /*KLK3* gene regulation. Previous studies have shown that androgen can promote cancer cell growth, indirectly regulate the TME, and induce DNA damage in prostate cancer [60]. It is worth noting that in C7 (Fig. 3f), many of the significant pathways were related to immunity or cell migration, further indicating the functional association with TLS region.

Based on these results, we focused on the prostaglandin-related immunological processes in cancer. Prostaglandins are involved in the regulation of immune responses and the enhancement of inflammation[61], and COX-1 and COX-2 are crucial enzymes in prostaglandin synthesis. The results demonstrated that *PTGS1*, the gene encoding COX-1, was highly expressed in C7, whereas *PTGS2*, the gene encoding COX-2, was highly expressed in both immune and cancerous regions (Fig. 3e and Supplementary Fig. S2). This may be related to the different functions of these two enzymes. COX-1 is a housekeeping gene that maintains basic prostaglandin levels [62], whereas Prostaglandin E2 (PGE_2_) induced by COX-2 may lead to tumor-promoting inflammatory responses, causing tumor growth and metastasis [63]. Furthermore, mPGES-1, mPGES-2, and cPGES are enzymes involved in PGE_2_ synthesis. mPGES-1 is encoded by *PTGES*, whereas mPGES-2 and cPGES are encoded by *PTGES2* and *PTGES3*, respectively. It can be observed that PTGES was significantly highly expressed in the C7, whereas *PTGES2* and *PTGES3* tended to have relatively higher expression in the cancerous regions. Previous studies have shown that mPGES-1 is an inducible isomerase, whereas mPGES-2 and cPGES are constitutively expressed [64]. We speculated that the abundance of *PTGES* in the C7 region induces a high expression of mPGES-1, majorly enhancing PGE_2_-mediated tumor-promoting inflammation. On the other hand, mPGES-2 and cPGES may be more inclined to maintain the synthesis of PGE_2_ in cancer tissues to sustain tumor growth and survival. It is worth noting that similar expression trends of related genes were observed by [65] for another cancer through patient-wise analysis, whereas we employed the SRT data with STForte analysis to refine the description of TME for prostate adenocarcinoma. Additionally, we analyzed the spatial interactions of prostaglandin-related genes recorded in CellPhoneDB [66] based on STForte’s results and COMMOT [36] (Fig. 3g, Supplementary Fig. S4, 5). The *PTGES* and *PTGES2* genes both influence the PGE_2_ receptors at non-tumor regions, including *PTGER2*, *PTGER3*, and *PTGER4*, with the invasive carcinoma or C7 regions as the starting points or important mediators in the TME. Activation of these PGE_2_ receptors has been shown to promote tumor spread through inflammation in various cancers [67]. These results illustrate the process of tumor infiltration via the C7 region caused by tumor-promoting inflammation, with PGE_2_ synthesis being one of the crucial processes involved.

### 2.5 Spatial smoothing control and fine-grained gene padding with oral squamous cell cancer

STForte is designed to address the over-smoothing issue typically associated with the massage-passing scheme in spatial domain identification by finetuning the cross-reconstruction factor *λ*_cross_. A detailed explanation of this mechanism was outlined in the methods section. We performed spatial domain identification on 12 oral squamous cell carcinoma (OSCC) samples from the 10x Visium platform with STForte to substantiate its applicability [68]. The ATTR encodings, generated under different *λ*_cross_ values, were clustered using the Louvain algorithm at a resolution of 0.4, followed by a manual consolidation of clusters based on histological images (Supplementary Fig. S6,7). An example result is presented in Fig. 4a. Upon decreasing *λ*_cross_ from 10 to 1, we observed increasingly refined identification of cancerous regions. Notably, when *λ*_cross_ was set to 1, STForte exhibited the potential to distinguish discrete punctiform cancer areas that were highly consistent with the pathologist’s annotations (Fig. 4a left 1-4). A sufficiently small *λ*_cross_ could encourage STForte to encode fine-grained, heterogeneous identities within spatial domains, which clustering algorithms can then uncover. The original Louvain clusters in azure and purple tended to provide suspicious separation of cancer core and leading edge (Fig. 4a right 1), where these findings were corroborated by the analysis of the OSCC source article [68].

**Fig. 4.**
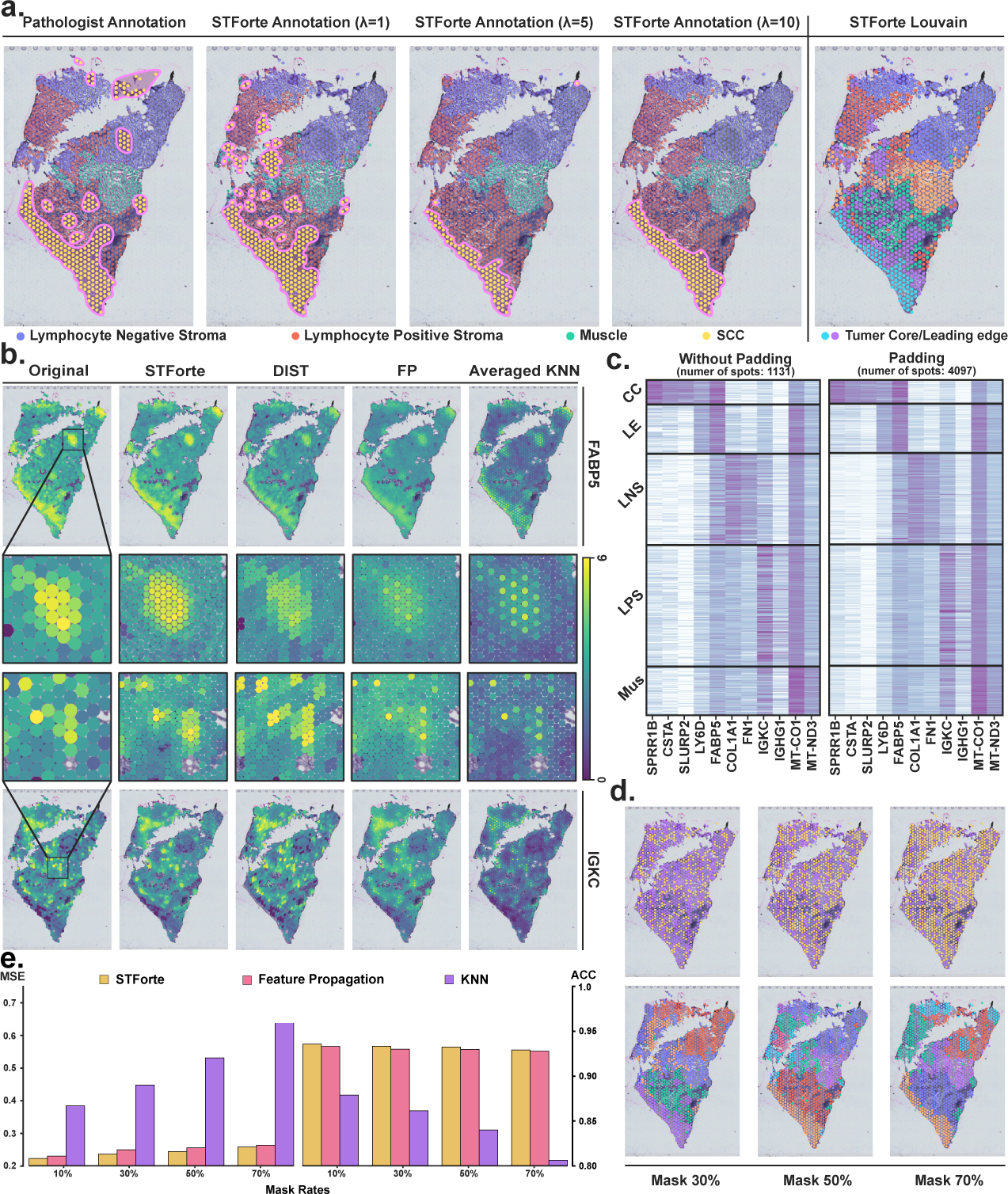
Example results of cancer edge identification with adjustment of λcross and fine-grained gene expression padding. **a,** The manually combined cluster labels (ATTR encoding) under different *λ*cross values (left 1-4). Original Louvain clusters when *λ*cross = 1 (right 1). **b,** Visualization of the comparison between padded gene expression of STForte and other methods for cancer leading edge (LE) marker *FABP5* and lymphocyte positive stroma (LPS) marker *IGKC*. **c,** Gene expression profiles of top highly variable genes in different tissue areas before and after gene padding. **d,** Robustness of spatial domain padding with randomly masked spots. **e,** Robustness of gene padding with randomly masked spots.

STForte excels at refining padding tasks at the gene expression level, rather than simply padding spatial domains. This is accomplished through semisupervised forecasting using XGBoost regression [69], leveraging the latent encodings generated by STForte. The expression patterns of highly variable genes, once padded by STForte, align with their original low-resolution counterparts across various tissue regions, as illustrated in Fig. 4c. Fig. 4b. provides the expression padding results of gene *FABP5* and *IGKC*, which are markers of cancer-leading edge and lymphocyte-positive stroma, respectively. To demonstrate the advantage of STForte, we further compared its performance with another image-free expression enhancement method DIST as well as two traditional methods: FP and averaged KNN. STForte was found to accurately deduce local expression profiles, even in scenarios where only sparse high-expression spots were detected. In contrast, the other methods produced noticeable artifacts. The padding results from DIST appeared to replicate and shift the original spatial profile, creating diamond-shaped patches that contradicted the expected local expression patterns. Both FP and averaged KNN underestimated the expression levels of inferred spots, leading to non-smooth padding outcomes. Besides, despite the domain identification results in Fig. 4a (right panel), where the area highlighted in Fig. 4b (gene *FABP5*) was not successfully categorized as a region of cancer leading edge, the gene padding results based on STForte’s encodings accurately recovered the expression pattern of its marker genes. This underscored the quality of STForte’s encodings but highlighted the necessity for a reasonable selection of clustering methods, especially for graph-based embedding methods.

To further assess the capabilities of STForte, we conducted an experiment by randomly masking spots within the original resolution and subsequently comparing the recovery outcomes using STForte with those using FP and the averaged KNN. The super-resolution approach utilized by DIST, which was derived from computer vision techniques, was not included in this evaluation, as it was not suitable for inferring the expression profiles of irregularly shaped spots. Fig. 4d highlights the masked spots in yellow and presents the clustering results generated by STForte beneath it. The findings indicated that STForte could accurately recover spatial details when the masked rate was below 30%. Even with a masked rate exceeding 70%, STForte was still able to recover spatial domain information with acceptable accuracy. This fully demonstrated the robustness and inferential ability of STForte. As the masked rate increased, the spatial information recovered became less detailed, which aligned with the observed downward trend in mean squared error (MSE) and accuracy (ACC) scores for all methods, as depicted in Fig. 4e. Notably, STForte achieved the lowest MSE and the highest ACC score among the evaluated methods.

### 2.6 Employing STForte for analysis of human dorsolateral prefrontal cortex (DLPFC) data

Next, we performed a systematic analysis using STForte on the human dorsal lateral prefrontal cortex (DLPFC) data generated by the 10x Visium platform. Clustering was conducted on the three latent encodings produced by STForte across 12 slices in the dataset to obtain spatial regions. Extended Data Fig. 5a shows the Adjusted Rand Index (ARI) and Normalized Mutual Information (NMI) between the clustering results from the different latent encodings and manual annotations. It can be observed that both ATTR and COMB encodings performed better in spatial region identification, with only minor differences between them (p-value *≫* 0.05 for Wilcoxon signed-rank test on ARI and NMI; details are provided in Supplementary Table S3). In contrast, TOPO encoding was relatively deficient in identifying regions that closely matched the manual annotations. We selected COMB encoding as the primary encoding in this investigation and compared STForte with other spatial region identification or latent embedding methods. Fig. 5a and Supplementary Fig. S8 illustrate the ARI, NMI, and identified spatial regions across 12 slices for STForte and other methods. STAGATE, GraphST, and STForte exhibited considerable performance (Wilcoxon signed-rank test results are provided in Supplementary Table S4). Methods that incorporate spatial information, including BayesSpace, DeepST, SpaGCN, and the three aforementioned approaches, exhibit a significant improvements in ARI and NMI compared to non-spatial methods. Furthermore, we focused on slice No. 151673 from the DLPFC dataset. Fig. 5b and Extended Data Fig. 5b illustrate the spatial domain identification results from the different methods and the three STForte encodings on slice No. 151673, respectively. Compared with Scanpy and scVI, nearly all spatial information-based methods could identify consistent and structured spatial regions within DLPFC tissue. It is worth noting that CA, despite being a non-spatial method, also identified structured and continuous spatial regions. The performance metrics in Fig. 5a also reveal that CA exhibited high ARI and NMI values for certain slices. This may be due to CA’s consideration of Pearson correlation coefficients between different spots, indirectly reflecting the spatial correlations among spots within those slices. We also conducted a trajectory analysis based on PAGA using latent encodings provided by different approaches (Extended Data Fig. 5c). The latent encodings of STForte and STAGATE presented fluent transitions from the white matter (WM) to Layer 1 (L1).

**Fig. 5.**
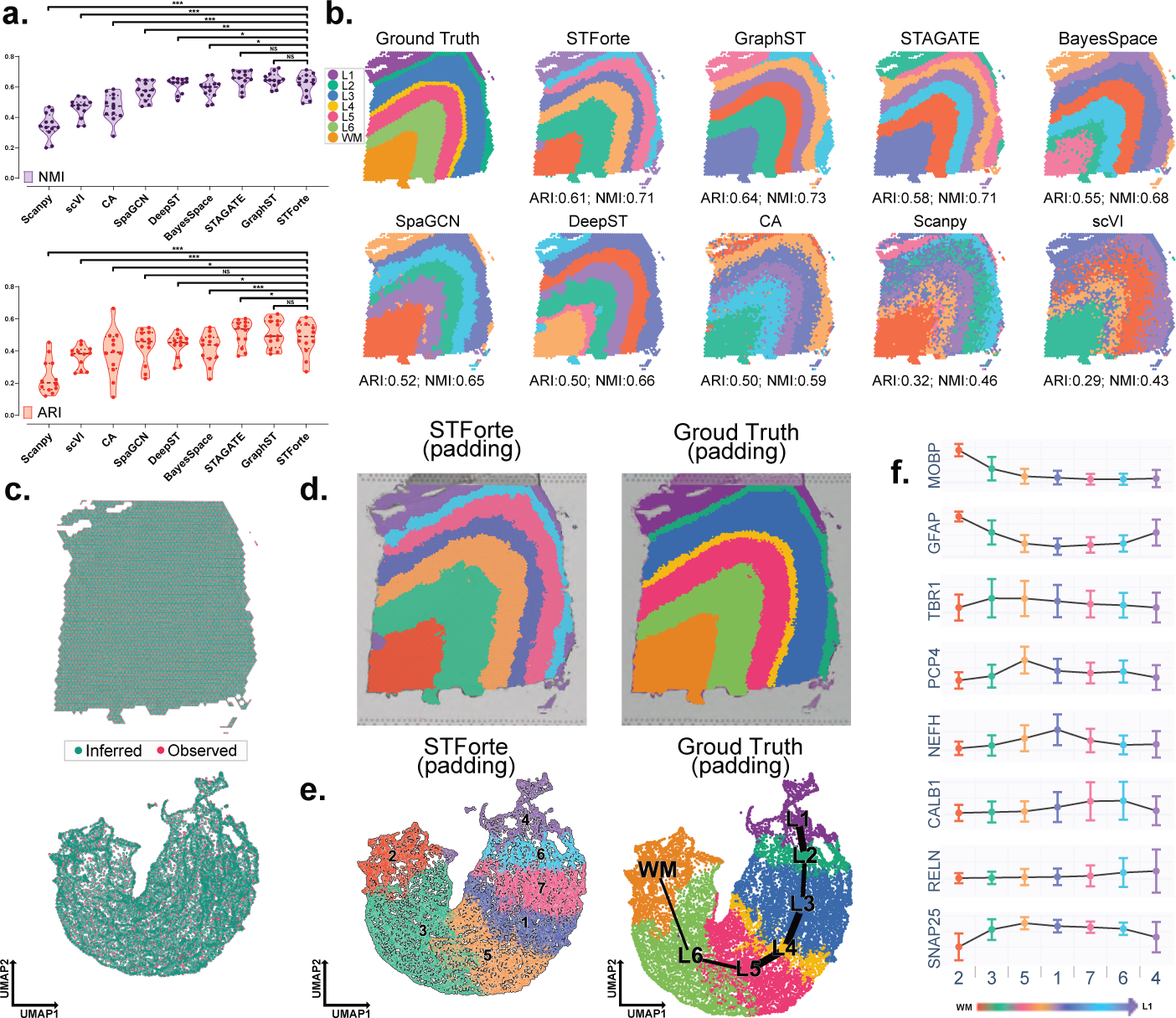
Investigations on the 10x Visium DLPFC dataset. **a,** Violin plot of spatial region identification performance across the 12 slices of the dataset using different approaches, quantified by the ARI and NMI metrics. The dashed lines represent the quartiles and median across 12 points. Asterisks denote significance levels: *: p *≤* 0.1; **: p *≤* 0.01; ns: p *>* 0.1. **b,** Spatial regions and performance metrics obtained by different approaches on slice No. 151673. The ground truth is derived from the manual annotations within the dataset. **c,** Spatial (top) and UMAP visualization based on the COMB encoding (bottom) show the spot instances, including observed spots (Observed) and unmeasured intervals (Inferred), obtained by STForte’s padding strategy for slice No. 151673. **d,** Spatial visualizations of the propagated annotations of the padding scenario using spatial identification results (left) or manual annotation (right) based on TOPO encoding. **e,** UMAP visualization based on the COMB encoding for the propagated annotations. Trajectory analysis was also carried out based on the propagated manual annotation and visualized in the UMAP space. **f,** Trend plots for the averaged expression levels of layer-specific marker genes based on the STForte spatial region identification are presented. The regions are ordered sequentially from WM to L1, with each error bar describing the standard deviation of expression level.

Moreover, STForte enabled fine-grained analysis, revealing the DLPFC structure through padding strategy (Fig. 5c). On slice No. 151673, all three STForte encodings provided continuous and consistent attribute imputations for unobserved locations (Extended Data Fig. 5d), allowing the consistent propagation of clustering results or manual annotations to unobserved locations to obtain more fine-grained spatial regions (Fig. 5d). Trajectory analysis incorporating the imputed properties in the padding scenario also demonstrated the continuity of the hierarchical structure from WM to L1 based on the COMB encoding (Fig. 5e). Additionally, we designed an RGB analysis to visualize the differences between the three encodings when applying the padding strategy to this dataset. Each encoding was processed using UMAP to reduce its dimensionality to three, followed by the standardization of each dimension and mapping to the RGB space. The results were visualized using spatial coordinates, as shown in Extended Data Fig. 5e. It can be observed that the encodings obtained through feature propagation using ATTR (ATTR FP) are not as smooth as TOPO or COMB encodings in the slice coordinate space, consistent with findings from other 10x Visium-based analyses discussed in the previous sections.

We further analyzed the expression patterns of layer-specific marker genes based on the results obtained from STForte. Specifically, *MOBP* is highly expressed in the white matter [70]. *GFAP*, a specific marker for astrocytes, is relatively enriched in the white matter and L1 of humans and primates [71, 72]. *TBR1*, *PCP4*, *NEFH*, *CALB*, and *RELN* are highly expressed at L6 [73], L5 [74], L3 [75], L2 [76], and L1 [77], respectively. *SNAP25*, a neuronal marker, is relatively enriched in the gray matter. However, due to its primary role in integrating and modulating information transmission, L1 contains a lower proportion of neurons compared to other grey matter layers [78]. Based on the spatial regions identified by STForte, these marker genes displayed distinct changes in expression levels, corresponding to the genuine structure of the human DLPFC (Fig. 5f). Furthermore, by leveraging the padding strategy, STForte can impute the gene expression levels for unobserved locations, yielding more fine-grained measurements of expression profiles (Supplementary Fig. S9).

Moreover, we utilized data from DLPFC No.151563-151676 for analysis in a multi-slice integration scenario. Specifically, we first aligned the spatial coordinates of different slices using the PASTE2 [79]. Based on the aligned coordinates, STForte was employed to construct an integrated latent space in a multi-slice perspective and conduct downstream analysis. The spatial region identification results in the multi-slice context are shown in Extended Data Fig. 6a. Notably, STForte yielded continuous and consistent spatial regions in the multi-slice view. Additionally, by combining the multi-slice spatial region analysis results from STForte with the paddings scenario of each individual slice for annotation propagation, we obtained more fine-grained spatial regions that incorporate unobserved locations.

Comparing the results with different methods (including STAGATE, CA, and GraphST) combined with PASTE2 analysis, STForte demonstrated the highest averaged ARI and NMI in multi-slice integration for the No.151563-151676 dataset. Regarding integration performance, STForte achieved the second-highest iLISI [80] (Extended Data Fig. 6b), surpassed only by GraphST. UMAP visualization also illustrated that STForte’s latent space could provide clear region delineation in the multi-slice scenario, as well as good cross-slice integration (Extended Data Fig. 6c).

### 2.7 Elucidating anatomical structures in Xenium mouse brain data using STForte

In this section, we show the capability of STForte in analyzing in-situ SRT datasets. Specifically, STForte was applied to the 10x Xenium mouse coronal brain dataset, which contains spatial measurements of over 130,000 cells in the entire mouse brain. Fig. 6a illustrates the referenced anatomical structure of the mouse coronal brain from the Allen Atlas [81]. The COMB encoding of STForte was employed for downstream spatial region identification via clustering, with the spatial and UMAP embedding results presented in Fig. 6b. It can be observed that different anatomical regions of the mouse brain were clearly identified, while the UMAP embedding space well depicted the functional distinction among cells within the entire brain. The clustering results were further annotated according to their spatial and anatomical characteristics, as summarized in Fig. 6c and Supplementary Table S5. The annotated domains, including the cerebral cortex, cerebral nuclei, hippocampus, thalamus, hypothalamus, ventricular system, and fiber tracts, aligned well with the reference anatomical structures shown in Fig. 6a. These domains are also distinctly separated in the UMAP space. Notably, the meninges and fiber tracts intersected with other regions in the center of the UMAP space. This might be due to the protective role of the meninges, leading to spatial proximity to other regions, while fiber tracts connect diverse neurons and neural structures, facilitating the association between various domains within the brain[82–84].

**Fig. 6.**
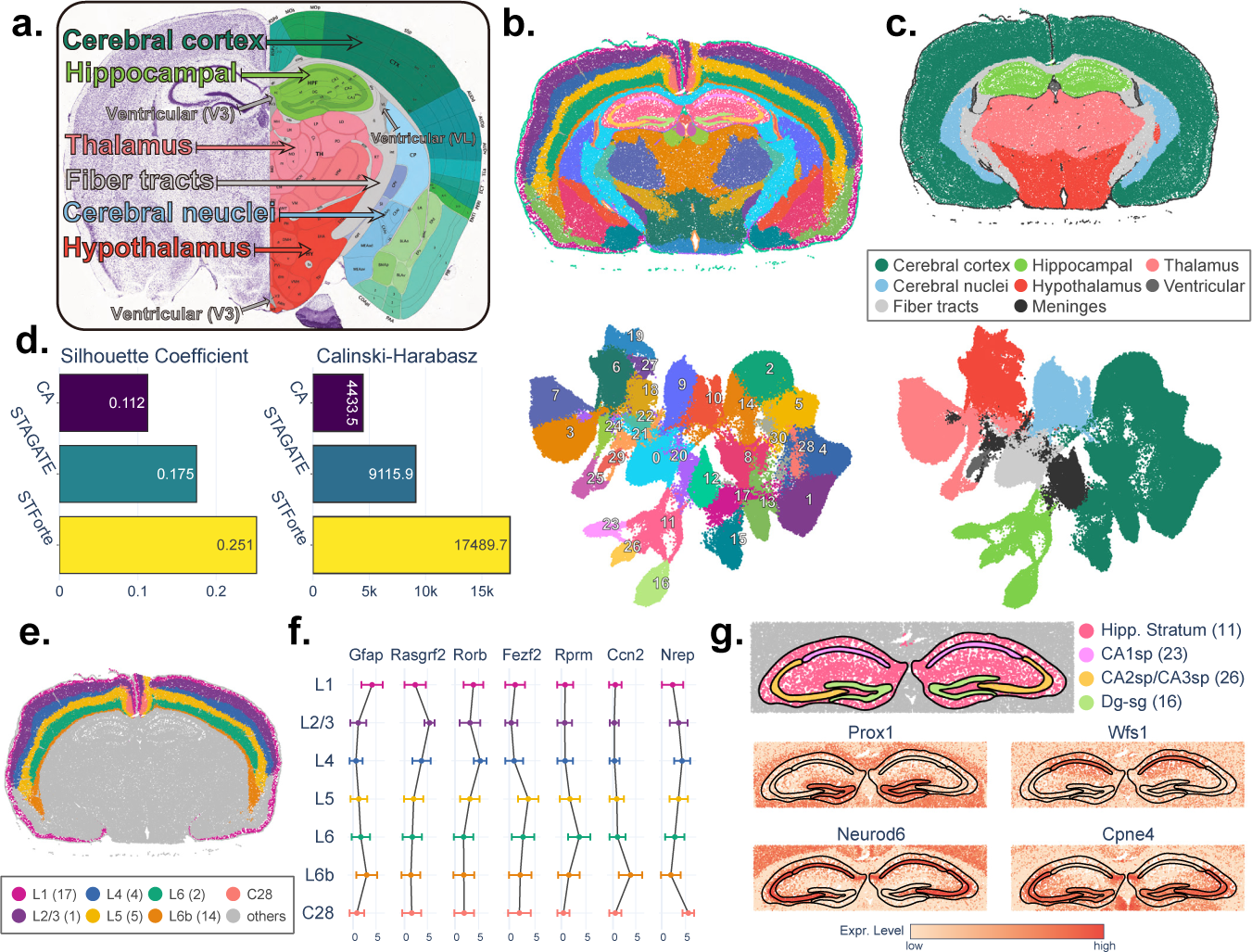
Analysis of Xenium mouse coronal brain data using STForte. **a,** Anatomical annotation of a coronal section of the mouse brain. Adapted from the Allen Brain Atlas. **b,** Spatial region identification based on STForte ATTR encoding on the Xenium mouse coronal brain dataset (top) and corresponding UMAP visualization (bottom). **c,** Spatial (top) and UMAP (bottom) visualizations of anatomical parcellation based on the summarized STForte results. **d,** Comparison of clustering performance metrics for different methods on this dataset. **e,** Visualization of the spatial regions identified by STForte in the Isocortex. **f,** Average expression levels of layer-specific marker genes in outer cortical layers. Error bars represent the standard deviation. **g,** Hippocampal regions identified by STForte and the spatial expression patterns of relevant marker genes.

We also performed STAGATE and CA workflows and compared them with those of STForte. Fig. 6d presents the comparison of clustering performance metrics across different methods, revealing that STForte achieves higher silhouette coefficients and the C-H index than the other two methods. Extended Data Fig. 7 illustrates the spatial regions and the UMAP results for each method. STAGATE also distinguishes the anatomical structures of the whole mouse brain, while CA, which does not consider spatial information, tends to differentiate cell characteristics. UMAP visualizations demonstrate that STForte’s latent space exhibits better separation between different anatomical domains than the other two methods.

Furthermore, we inspected the expression patterns of known marker genes to evaluate whether STForte accurately delineates subregions in different anatomical domains. Fig. 6e,f illustrate the regions of the Isocortex identified by STForte and the expression levels of different layer-specific genes within these divisions (Wilcoxon rank-sum tests result is provided in Supplementary Table S6). Specifically, *Gfap* showed high expression levels in layer 1 (L1) of the Isocortex[85, 86], while *Rasgrf2* was highly expressed in layer 2/3 (L2/3)[87]. *Rorb* exhibits strong expression in layer 4 (L4)[88], and *Fezf2* was highly expressed in layer 5 (L5)[89]. *Rprm* was highly expressed in Layer 6 (L6)[90], and *Ccn2* was strongly expressed in Layer 6b (L6b)[91]. These layer-specific gene expression patterns highlight the distinct genetic profiles of each cortical layer, and demonstrate the ability of STForte to accurately identify these layers based on their unique molecular signatures. Moreover, cluster 28 in the Isocortex was identified separately, and *Nrep* exhibited a higher expression level in this region. This region may correspond to the retrosplenial cortex (RSC) portion of the mouse brain, and the RSC is considered to play a significant role in memory and cognitive processes[92]. The expression of *Nrep* is associated with neural regeneration and cognitive function to a certain extent[93, 94]. We also observed gene expression patterns in the hippocampus (Fig. 6g and Supplementary Table S7). STForte divided the hippocampal region into four parts, corresponding to the hippocampal stratum (Cluster 11), CA1sp (Cluster 23), CA2sp and CA3sp (Cluster 26), and the dentate gyrus (Dg-sg, Cluster 16). It can be seen that *Prox1* was relatively enriched in Dg-sg[95], and *Wfs1* was relatively enriched in CA1sp[96]. *Neurod6* was highly expressed throughout the entire Ammon’s Horn (CA) region[95, 97], while *Cpne4* had a relatively higher expression level in CA2sp/CA3sp and Dg-sg[98]. These results demonstrated that STForte can perform precise anatomical region identification of complex brain structures at single-cell resolution based on 10x Xenium data.

## 3 Discussion

Extracting expressive encodings from single-cell and spatial transcriptomic data forms the foundation for various downstream analysis tasks. Enhanced spatial resolution could further benefit the analysis to achieve more finegrained results. In this study, we have presented a representation learning and spatial enhancement method STForte for SRT data analysis. STForte separately considers the expression and spatial information via a pairwise auto-encoder framework, and it can generate the encodings at an enhanced resolution merely based on the connectivity of the spatial graph. STForte’s spatial enhancement strategy explicitly incorporates neighboring information, resulting in better spatial continuity compared to existing studies. Additionally, STForte naturally achieves spatial enhancement at both the spatial domain level and the gene-specific level. For assigning unobserved spots, STForte is compatible with customized spatial locations, making it suitable for more flexible tasks, including local spatial enhancement and local expression reconstruction.

Introducing the graph structure during the encoding process has been a popular solution to incorporate spatial information. These approaches usually implicitly assume that neighboring spots share similar expression profiles and tend to preserve the spatial correlation among neighboring spots to a certain extent when extracting latent encodings. Appropriate consideration of the spatial graph structure has proven successful in extracting representative encodings. However, excessive preservation of the spatial graph structure in encoding can be counterproductive, particularly with data from nonconsecutive spatial domains. To avoid over-emphasizing the reconstruction of the spatial graph structure, STForte employs a heuristic cross-reconstruction training strategy and manually regulates the balance between the attribute and topology encoders in incorporating different types of information. By integrating prior knowledge of the SRT data and its corresponding histological image, STForte effectively addresses the aforementioned issues and accurately describes the complex and delicate tumor micro-environment. To further demonstrate this advantage of STForte, we compared its performance with the typical non-spatial aware method CA and one of the current graphbased representation learning benchmarks STAGATE. Results supported the effectiveness of STForte’s training scheme and exhibited a certain degree of controllability.

STForte could be applied to data generated by different SRT technologies, including but not limited to Visium, Xenium, and Stereo-seq, which are the technologies shown in this study. Moreover, STForte carries special optimization for large SRT data regarding the sparsity of spatial graphs. It could accomplish exhaustive training on SRT data of approximately 0.8 million spots without spatial enhancement within 1 hour on a single RTX TITAN or RTX 3090. Applying analysis with local spatial enhancement by adding a few unobserved spots may not significantly prolong the training time. Global spatial enhancement is unnecessary since data of this scale is usually of single-cell or sub-cellular resolution. For SRT data of the DLPFC 151673 scale (*∼*3500 spots), the training time of STForte with spatial enhancement (*∼*14,000 spots) is approximately 2 minutes on the same GPUs mentioned above. Compared to existing spatial enhancement approaches, STForte demonstrates a notable efficiency advantage but is still less efficient than specialized representation learning methods.

SRT data exhibit significant diversity and potential heterogeneity, influenced by factors such as disease type, sample area, sampling and embedding methods, and sequencing technologies. It is challenging for a single representation method to adapt to all possible SRT data. This study aims to offer an improved representation learning method combined with a flexible spatial enhancement approach and highlights the need to consider the potential challenges of applying graph-based algorithms in SRT analysis. Previous studies have provided several tricks for addressing the oversmoothing problem via replacing the outer-product decoder of the vanilla graph autoencoder with an expression reconstruction decoder and applying PCA on the latent encodings. These approaches focus on losing spatial reconstruction constraints and dropping the potential graph structure contained in the low-frequency domain of the normalized Laplacian matrix via its similarity to the covariance matrix of encodings. However, they still lack control of spatial effects and cause extra information loss. Heterogeneous graph and graph structure learning (GSL) could provide better performance for addressing this issue but may require significant computational resources. Therefore, future studies could focus on constructing representation learning methods that consider the potential problems caused by spatial graph structure with superior efficiency and performance.

## 4 Methods

### 4.1 Problem statement

STForte constructs a graph 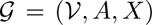 from the spatial coordinates and expression matrix of the SRT data. Specifically, 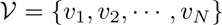 denotes the node set that represents the total *N* spots/cells. Spatial coordinates are used to establish the spatial topology of different spots/cells, denoted as an adjacency matrix *A ∈ {*0, 1*}^N×N^*, which is constructed using either a distance-based neighboring strategy or a k-nearest neighbor (KNN) strategy, determined by whether the SRT protocol is formulated with a regular lattice (see Extended Data Fig. 1a and Supplementary Note S1 for more details). Expression profiles, denoted by 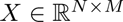, can be either a matrix of gene expression values or results from a non-spatial dimensionality reduction approach (e.g., PCA).

STForte is designed to handle spots/cells of unobserved locations without given expression profiles. To this end, the node set *V* consists of two subsets: 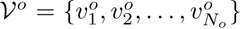 representing the *N_o_* observed spots/cells, and 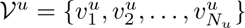 representing the *N_u_* unobserved spots/cells. Note that 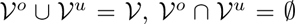, and *N_o_* + *N_u_* = *N*. Subsequently, the adjacency matrix and expression profiles corresponding to the observed spots/cells are represented as 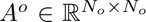 *^o^* and 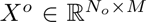, respectively. The expression profiles of unobserved locations, denoted by 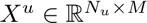, are not measurable and thus missing.

The objective of the STForte architecture is to extract graph information (*G*), while recovering node entities of unobserved locations (*V^u^*). Besides, the STForte model aims to comprehensively consider the spatial topology (*A*) and expression profile characteristics (*X*) to provide a contextual description of SRT data. Inspired by Chen et al.[99], STForte employs a pairwise autoencoder architecture to derive *l*-dimensional latent-space representations that encompass spatial topology and expression-specific information. As a result, the latent encodings derived by STForte facilitate downstream analysis for both homogeneous and heterogeneous SRT data as well as enable the inference of biological patterns at unobserved locations, thus providing additional analytical capabilities.

### 4.2 Latent Encoding Framework

The latent encodings of SRT data are obtained through a pairwise graph autoencoder (Extended Data Fig. 1b), where gene expression profiles and spatial topology information are processed by two distinct encoding structures:

i. The attribute encoder, a multi-layer perceptron (MLP) 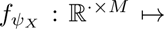 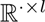*^l^*, maps expression profiles to *l*-dimensional latent variables, with *ψ_X_* denoting the attribute encoder parameters.
ii. The topology encoder is represented 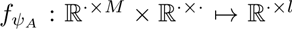 that maps the topology information of the graph converted from SRT data to *l*-dimensional latent variables through a Pathfinder Discovery Network [100] (Supplementary Note S2), where *ψ_A_* is the topology encoding parameters.

Subsequently, decoders are designed to handle the restoration from the latent-space representation to the expression profiles and topology information:

i. The attribute decoder part 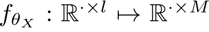 is an MLP that restores expression profiles from the latent variables, where *θ_X_* indicates the corresponding parameters.
ii. For the topology part, the latent variable is processed by the topology decoder defined as 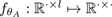*^·^* which is an MLP parameterized by *θ_A_* with an inner product operation followed by the last layer. The inner product operation is used to restore the adjacent matrix 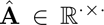 as 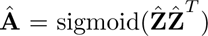, where 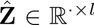 is the output from the last layer of the MLP.

### 4.3 Match between latent encodings

To encourage the integration of information between expression profiles and spatial topology, and to enable the imputation of unobserved locations for SRT data analysis, the latent encodings of both attributes and topology are aligned under the assumption that they are associated and can be projected into a shared latent space. Specifically, 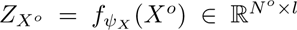 represents the attribute latent encoding of spots derived from the observed expression profiles. For topology encodings, a feature propagation method [101] is employed, leveraging the spatial topological structure and observed expression values to impute unavailable expression profiles at unobserved locations (Supplementary Note S3), with these imputed profiles denoted as 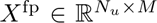. The topology encoding is computed as 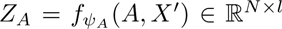, where 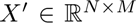 represents the expression profiles comprising *X^o^* and *X*^fp^. Consequently, two adaptive approaches are implemented to align the latent spaces of *Z_X_o* and *Z_A_* in STForte training strategy.

#### 4.3.1 Reconstruction matching

In the vanilla scheme of the autoencoding framework, the reconstruction loss compels the latent encodings to reproduce the observation [102]. Furthermore, in STForte, we assume that the attributes and topological features of each node within the graph are intrinsically correlated and can be projected into a shared latent space. Based on the assumption of shared latent space, decoupled latent encodings can be utilized to reconstruct their counterpart information. Consequently, the reconstruction of attribute information is formulated as follows:

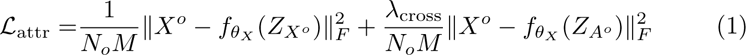

where *λ*_cross_ denotes the weight factor for the cross-information reconstruction trade-off and 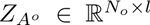 represents the topology latent encoding corresponding to the observed spots/cells 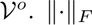 denotes the Frobenius norm. The first term in Equation 1 represents the mean square error for measuring the self-reconstruction quality of the attribute information, and the second term represents the cross-reconstruction loss used to restore the attribute information from topology latent encoding. Furthermore, the reconstruction of topology information is formulated as follows:

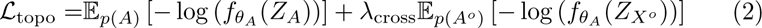

In Equation 2, the first cross-entropy (CE) term measures the quality of self-reconstruction for topology information, whereas the second CE term qualifies the cross-reconstruction of recovering the node neighboring relations from the observed attribute latent encoding. Subsequently, the overall reconstruction loss is formulated as follows:

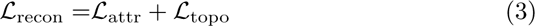

The objective of the reconstruction matching process is achieved by minimizing *L*_recon_ with respect to the parameters *ψ_X_, ψ_A_, θ_X_, θ_A_* during training. This allows latent encodings to learn condensed information from the original data, whereas the cross-reconstruction process enables information sharing between latent encodings.

#### 4.3.2 Adversarial distribution matching

To further impose closeness between the latent encodings of attribute and topology, STForte utilizes an adversarial-based approach [103] to match their latent spaces. Denote *Z^′^* as the latent samples drawn from a reference prior *q*(*Z*). The optimization refers to the minimax objective [104], which can be defined as follows:

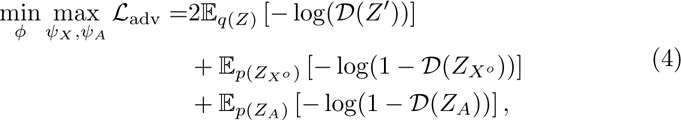

where *D*, parameterized by *ϕ*, is an MLP-based discriminator shared among different latent encodings. In STForte, the reference distribution is configured as a standard Gaussian distribution. Equation 4 conducts adversarial distribution matching to encourage the manifolds of topology and attribute latent encodings to be close to the reference prior, implicitly imposing the closeness between the attribute and topology features.

### 4.4 Overall objective

Combining the above loss functions and objectives, the main objective function of the STForte model in the training process is as follows:

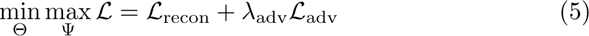

where Θ = *θ_X_, θ_A_, ψ_X_, ψ_A_, ϕ* and Ψ = *ψ_X_, ψ_A_*. The weight factor *λ*_adv_ is utilized for the trade-off between the reconstruction and adversarial distribution matching.

### 4.5 Encodings and property propagation

Extended Data Figure 8 illustrates the schematic descriptions of the STForte encodings. The trained model generates two encodings from the corresponding SRT data: attribute encoding (ATTR, *Z_X_*) to encode the expression pattern information in the observed spots, and topology encoding (TOPO, *Z_A_*) to compress the spatial patterns for both the observed spots and spots with unobserved transcripts. For unobserved locations, feature propagation is adopted to impute the ATTR encoding. In addition, ATTR and TOPO encodings are further concatenated to formulate combined encodings (COMB).

The generated encodings can be utilized for various downstream analyses of the SRT data. COMB encodings can be more effective for spatial region identification or related analyses of consecutive tissue domains (e.g., anatomical regions) by considering a trade-off between expression patterns and spatial connectivity. For the analyses of tissues with higher complexity and heterogeneity (e.g., tumor micro-environment), ATTR encoding is more suitable for identifying distinct patterns by mitigating the influence of spatial connectivity. In addition, the trade-off of spatial connectivity can also be handled implicitly through adjusting the cross-reconstruction factor *λ*_cross_.

To enable fine-grained analysis of SRT data, STForte includes additional utilities for propagating annotation or imputing gene expression levels for unobserved spots. Specifically, the latent encoding of the observed spots serves as training data to fit the XGBoost model [69]. Subsequently, the properties of unobserved spots are predicted using the fitted XGBoost. Annotation propagation is handled as a classification task, whereas gene imputation is a regression task used to predict the expression levels of specific genes. Normalization of gene expression levels included log-transformation and normalization according to library size from the Scanpy package [105]. Because TOPO encoding considers more consecutive information about spatial connectivity, it is adopted as encoding for propagation or imputation by default in analyses. Note that other encodings are also included in the investigation and evaluation of the encodings in the result sections.

### 4.6 Details on analysis

We ensured utmost uniformity in the preprocessing and the selection of hyperparameters across different datasets. However, slight variations may be present owing to differences in the SRT techniques and spatial heterogeneity. Here, we describe the details of analysis according to different investigated datasets as follows.

#### 4.6.1 10x Visium mouse olfactory bulb dataset

Lebrigand et al. [38] processed olfactory bulb tissue samples from C57BL/6 mice (*>*2 months old) using the 10x Genomics Visium protocol. The original dataset comprised 918 spots and 31,053 genes. Spatial neighbors were constructed based on the distance-based strategy, which is the default setting for 10x Visium datasets in STForte (Supplementary Note S1). Outlier cells expressing *<* 200 genes were masked as unobserved locations. Additionally, padding strategy was used to identify gaps within the original spots as unobserved locations. PCA with 300-dimensions was applied for the pre-dimensional reduction of the expression counts using “sc.pp.pca” from the Scanpy library. Subsequently, STForte was applied to the preprocessed data using default hyperparameters. For spatial region identification, “mclust”, with the number of clusters set to six, was applied to different STForte encodings. The primary investigations and visualizations were performed using COMB encoding for the observed spots, taking into account the spatial continuity of the laminar structure in the olfactory bulb.

#### 4.6.2 Stereo-seq mouse olfactory bulb dataset

The adult mouse olfactory bulb data obtained by Stereo-seq was binned to a resolution approximating the cellular level (*≈*14 *µ*m)[27]. We retained only on-tissue cells based on the corresponding cell quality indices [11]. Subsequently, cells with defective expression profiles were masked (i.e., identified as unobserved locations) according to the number of expressed genes for each cell and the provided cell quality indices. The resulting dataset contains 19,326 cells (including unobserved locations) and 22,780 genes. Considering that the number of cells exceeded 15k, PCA based on the PyTorch [106] backend was used to generate a 300-dimensional pre-dimensional reduction for the expression profiles. In addition, a k-nearest neighbor (KNN)-based strategy (*k* = 18) was employed to construct spatial neighbors, taking into account the non-lattice identity of the dataset. STForte was applied to the preprocessed data using default hyperparameters, except for the sparse manner in the reconstruction of the adjacency matrix, considering the size of the dataset. Consequently, Leiden clustering [107] (”sc.tl.leiden” from Scanpy) with a resolution of 0.3 was adopted for spatial region identification. The primary investigations and visualizations were proceeded with COMB encoding.

#### 4.6.3 Prostate adenocarcinoma dataset

The 10x Visium spatial formalin-fixed, paraffin-embedded (FFPE) sample of prostate adenocarcinoma was obtained from the 10x Genomics repository, which contains 4,371 spots and 17,943 genes. The preprocessing steps for the dataset were the same as those used for the 10x Visium mouse olfactory bulb data, with the only difference being the exclusion of outlier spot filtering due to the relatively intact expression profile quality across this sample. STForte was applied with *λ*_cross_ = 1 (default setting is 10) considering the heterogeneity of the tumor micro-environment (TME). Leiden clustering was employed for spatial region identification. We investigated the clustering results by varying the resolution from 0.1 to 1 with a step size of 0.05 and selected the best results based on the highest silhouette coefficient. Primary investigations and visualizations were performed using ATTR encoding, considering the heterogeneity of TME. Furthermore, we applied COMMOT [36] to investigate spatial cell-cell communications based on the STForte results.

#### 4.6.4 Human OSCC dataset

The OSCC dataset contained 12 frozen and optimal cutting temperature embedded tissue samples and spatial transcriptomics was performed using 10x Visium [68]. Each data sample contains 1200 spots and 16,000 genes. Genes that were not detected in any spot were removed during the preprocessing and then followed by dimensional reduction to 300 via PCA. Spatial enhancement preparation and graph construction were accomplished using KNN reconstruction with hyperparameter k=18. Then STForte was applied to the prepared data with the neurons in the two GNN layers reduced to 100 and 50 respectively, and *λ_cross_* = 1. Spatial domain identification was achieved through the Louvain algorithm based on the attribute encodings.

#### 4.6.5 Human DLPFC dataset

The famous six-layered human dorsolateral prefrontal cortex (DLPFC) dataset included 3 groups of samples (4 continuous sections per group) generated using the 10x Visium technique [108]. The pre-processing, dimensional reduction and graph construction schemes of the DLPFC dataset were the same as those dealing with the OSCC dataset. Two larger GNN layers of 200 and 50 neurons and *λ_cross_* = 10 were adapted during the training process. Spatial domain identification was performed over concatenated attribute and topology encodings using R package ‘mclust’. Cluster numbers were explicitly set to 7 for samples in group 1 (151507-151510) and group 3 (151673-151676), and 5 for samples in group 2 (151669-151672). For multi-slices analysis, PASTE-v2 [109] was first applied to align the 2D coordinates of different samples in one group, followed by the same pre-processing and dimensional reduction scheme. The cross-slices distance-based neighbor graph was constructed with d=145, then the STForte model with the same parameters as training on a single slice. Clustering adapted “mclust” package and spatial enhancement were done by extending the multi-slices clustering results over the topological encodings of each single-slice model.

#### 4.6.6 10x Xenium mouse brain dataset

10x Genomics obtained a 10*µ*m fresh frozen tissue section of the coronal brain from a C57BL/6 mouse, which was prepared following the 10x Xenium In Situ protocol [110]. The dataset is publicly available in the 10x Genomics repository and contains 130,870 cells and 248 genes. During preprocessing, cells with counts less than 10 were excluded using “sc.pp.filter cells” from the Scanpy library. Expression counts were normalized according to their total counts across all genes and log-transformed. The processed expression profiles were used as direct inputs for STForte. Spatial neighbors were constructed using the KNN-based strategy with *k* = 10. STForte was applied to the preprocessed data in a sparse manner, with 500 epochs for the training process. Subsequently, Leiden clustering with a resolution of 0.85 was employed for spatial region identification. Investigations and visualizations were performed using the COMB encoding.

#### 4.6.7 Evaluation metrics

Evaluation metrics used in this study is detailed in Supplementary Note S4. These metrics were selected to rigorously assess various aspects of our spatial transcriptomics analysis.

### 4.7 Compared methods

Descriptions of the methods selected for comparison are provided in Supplementary Note S5.

## Data availability

All datasets included in this study are available in raw data from their corresponding resources. The 10x Visium mouse olfactory bulb dataset is available from the Gene Expression Omnibus (GEO) repository (accession No.GSM4656181). The Stereo-seq mouse olfactory bulb dataset [8] is accessible on https://github.com/JinmiaoChenLab/SEDRanalyses/tree/master/data. The 10x Visium FFPE prostate adenocarcinoma dataset can be obtained from the 10x dataset repository (https://www.10xgenomics.com/datasets/human-prostate-cancer-adenocarcinoma-with-invasive-carcinoma-ffpe-1-standard-1-3-0). The human OSCC dataset is available from the GEO repository (accession No. GSE208253). The DLPFC dataset can be downloaded from https://research.libd.org/spatialLIBD/. The 10x Xenium dataset can be obtained from the 10x dataset repository (https://www.10xgenomics.com/datasets/fresh-frozen-mouse-brain-for-xenium-explorer-demo-1-standard, Full coronal section).

## Code availability

The STForte software code is implemented by Python and is public available at https://github.com/poncey/STForte.

## Acknowledgments

This work was financially supported by the Center for Intelligent Drug Systems and Smart Bio-devices (IDS2B) from The Featured Areas Research Center Program within the framework of the Higher Education Sprout Project and Yushan Young Fellow Program (113C51N055) by the Ministry of Education (MOE), National Science and Technology Council (NSTC 112-2740-B-400-005, 113-2321-B-A49-025, 113-2634-F-039-001, 113-2221-E-A49-160-MY3), and The National Health Research Institutes (NHRI-EX113-11320BI) in Taiwan.

---

**Extended Data Fig. 1.**
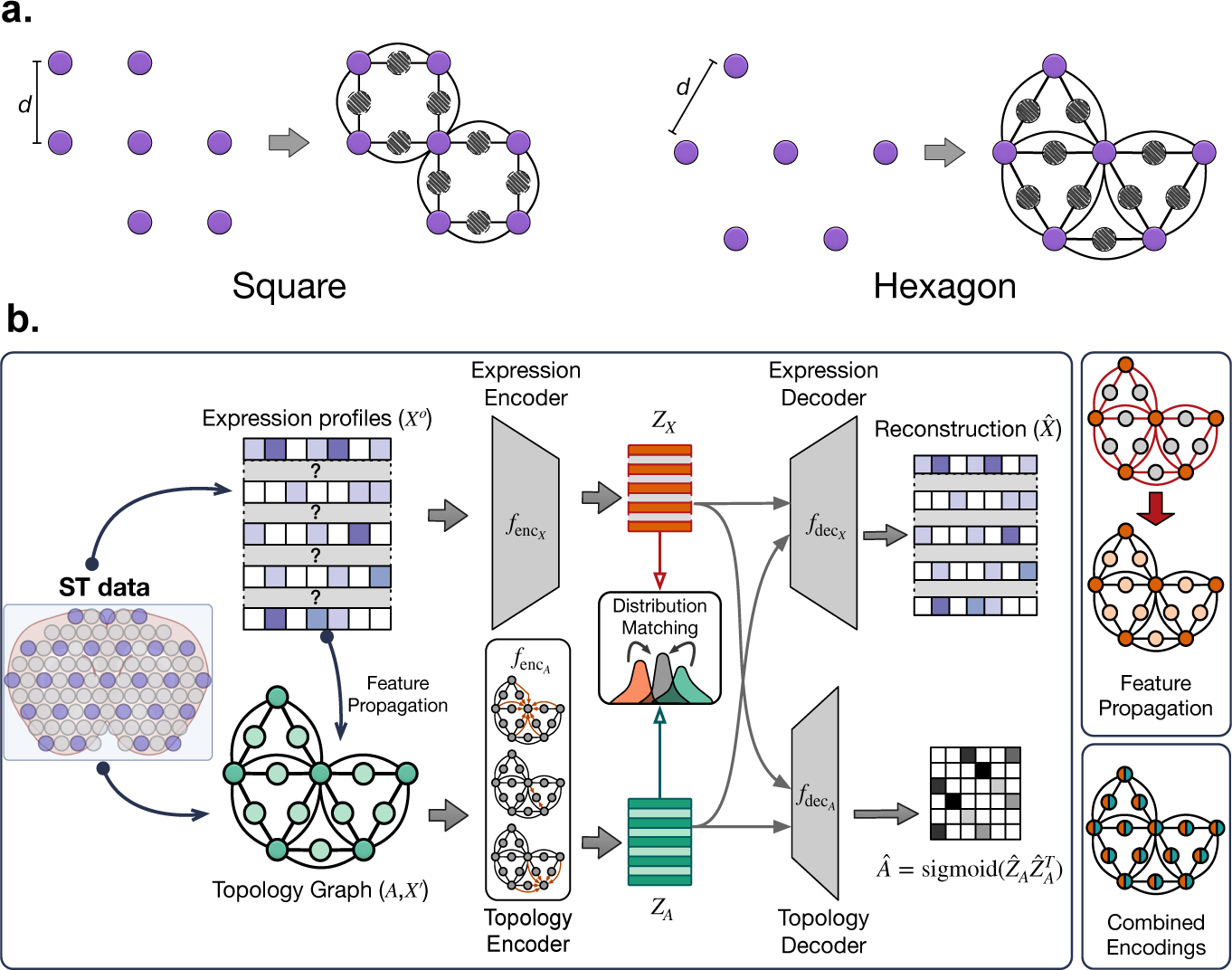
**a,** Spot padding strategy for intervals of unobserved expressions in regular-lattice SRT data. For square-arranged lattice data, such as Spatial Transcriptomics, or hexagon-arranged data, such as 10x Visium, new spots are inserted based on the center-to-center distance (*d*). **b,** STForte latent encoding framework. First, expression profiles and spatial topology are obtained from the SRT data. The extracted information is subsequently fed into a pairwise graph autoencoder (GAE), where the expression profiles of observed locations (*X°*) and topology graph, including spatial neighbors *A* and expression profiles with imputed data for unobserved locations thorugh feature propagation *X^′^*, are separately encoded into their corresponding latent spaces (*Z_X_*and *Z_A_*). The fitting process includes (1) self/cross-reconstruction streams to restore the expression attributes (*X*^^^) and adjacent matrix (*A*^^^) for spatial-aware dimensional reduction and latent-space alignment, and (2) adversarial distribution matching to further control the latent space of both encondings. Finally, *Z_X_*(ATTR), *Z_A_*(TOPO), and their combined encodings through concatenation (COMB) can be used for diverse downstream analyses.

**Extended Data Fig. 2.**
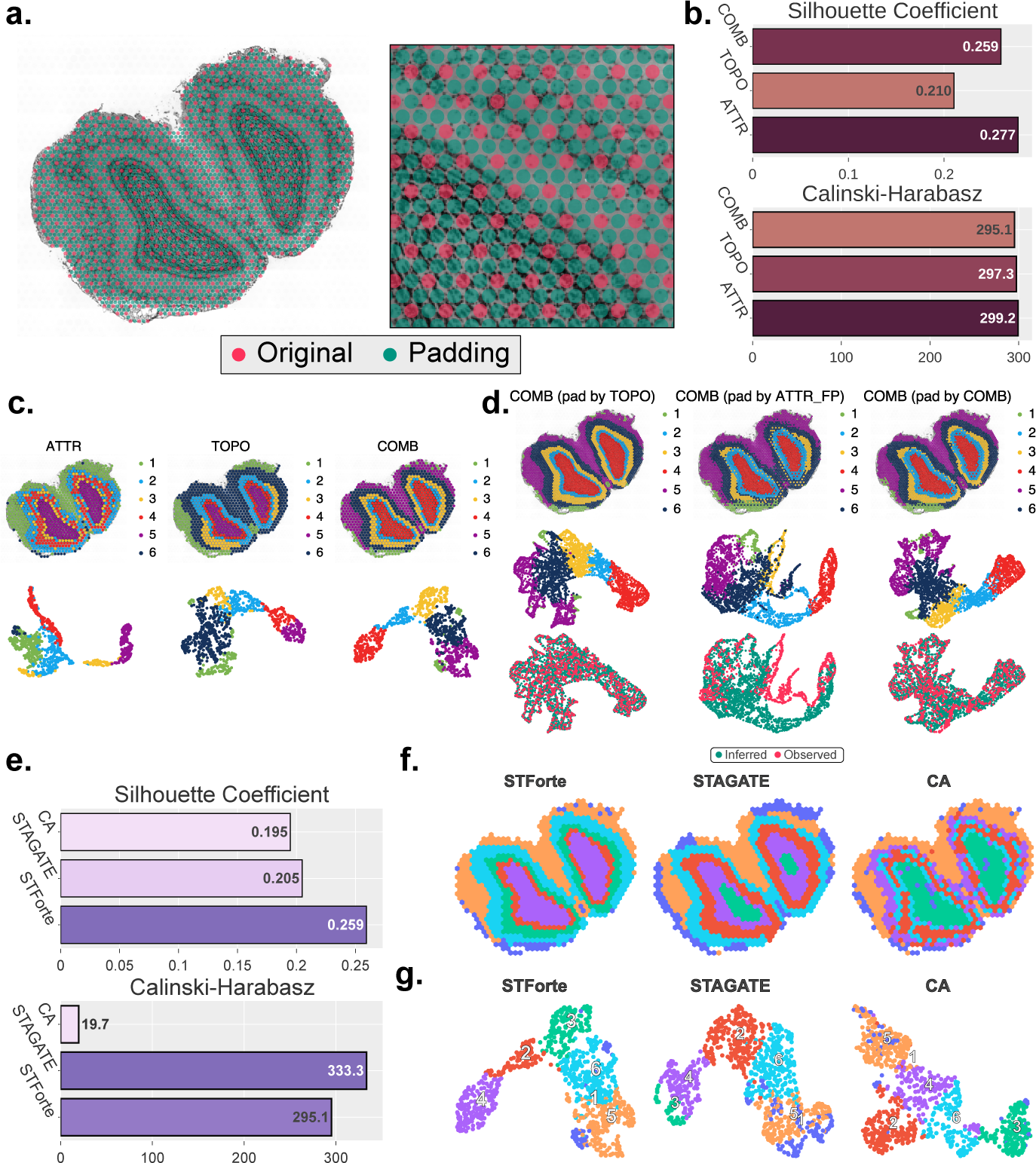
**a,** Spot instances including well-measured spots (original) and unmeasured intervals generated by STForte’s padding strategy. **b,** Comparison of clustering metrics among different STForte encodings, including the Silhouette Coefficient and the Calinski-Harabasz index. **c,** Spatial regions (top) and UMAP visualizations (bottom) of different STForte encodings. **d,** Spatial regions (top), UMAP visualizations based on region annotations (middle), and spot instances (bottom) of different STForte encodings under padding scenario. **e,** Comparison of clustering metrics for different dimensional reduction methods (STForte, STAGATE, and CA). **f,** Spatial regions identified by different methods. **g,** UMAP visualizations obtained by different methods.

**Extended Data Fig. 3.**
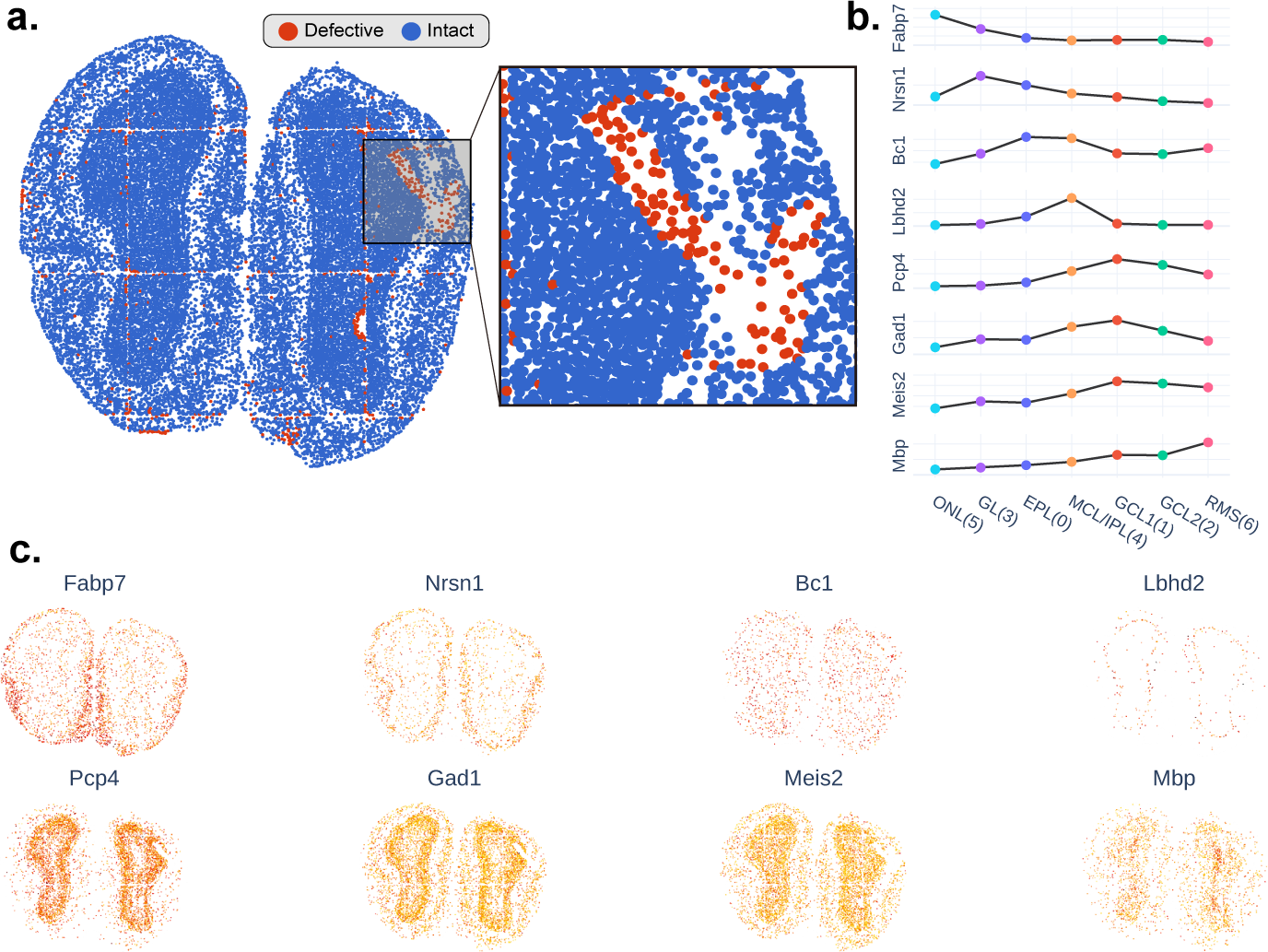
**a,** Spot instances including well-measured cells (Intact) and-low-quality cells (Defective) masked for the STForte process. **b,** The tendency-plot shows the mean expression of different genes within different spatial regions. **c,** Visualization of spatial expression levels for the layer-specific genes.

**Extended Data Fig. 4.**
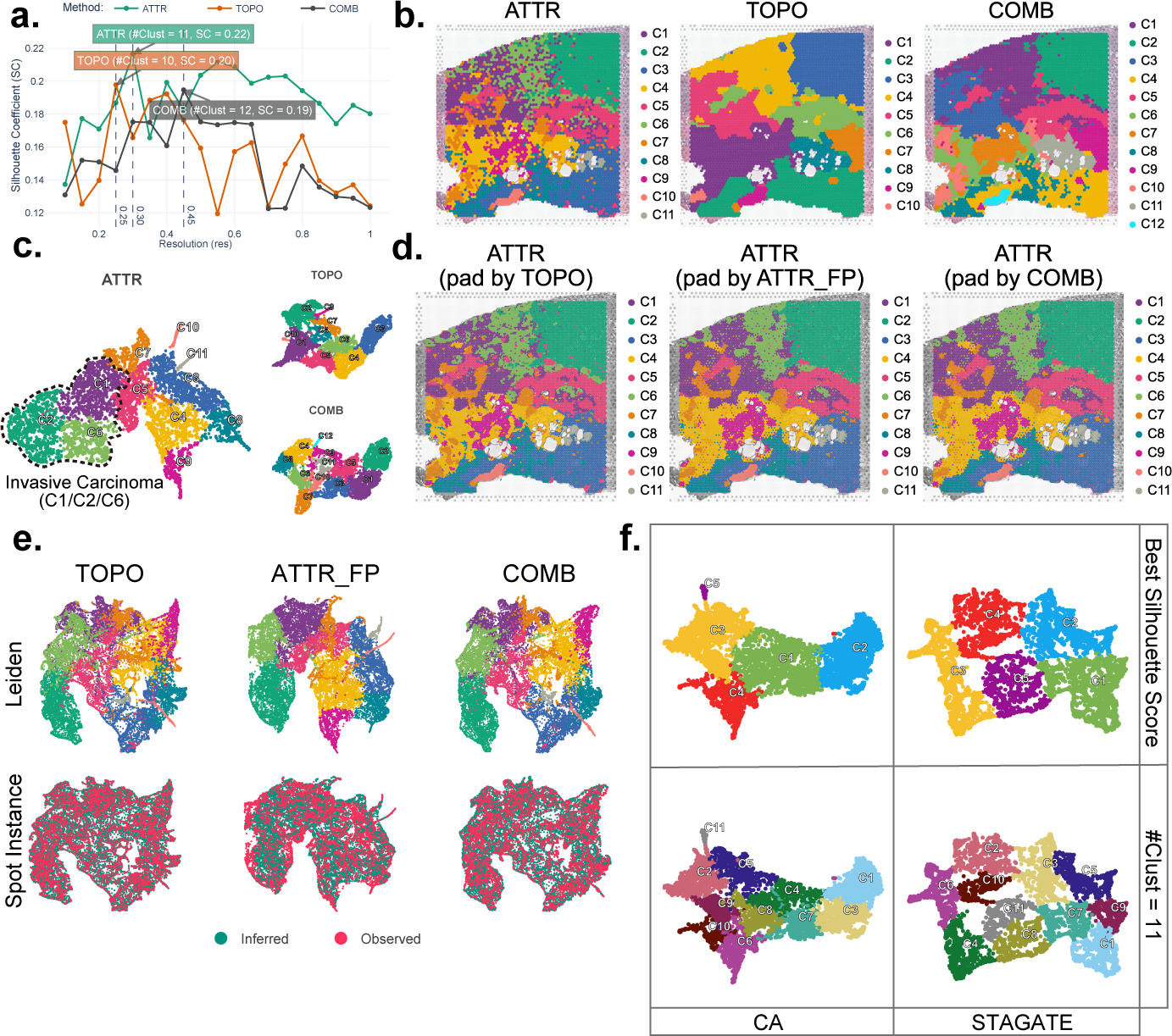
**a,** Silhouette coefficients (SC) of different STForte encodings under the Leiden method at various resolutions. #Clust indicates the number of clusters. **b,** Spatial regions of different encodings at the best silhouette coefficient under Leiden clustering. **c,** UMAP visualization of different STForte encodings. For ATTR, C1, C2, and C6 correspond to invasive carcinoma. **d,** Spatial regions of different STForte encodings under padding scenario. **e,** UMAP visualizations based on Leiden results and spot instances of different STForte encodings under padding scenario. **f,** UMAP visualization shows the Leiden results from STAGATE and CA at their respective best silhouette coefficients or when #Clust=11.

**Extended Data Fig. 5.**
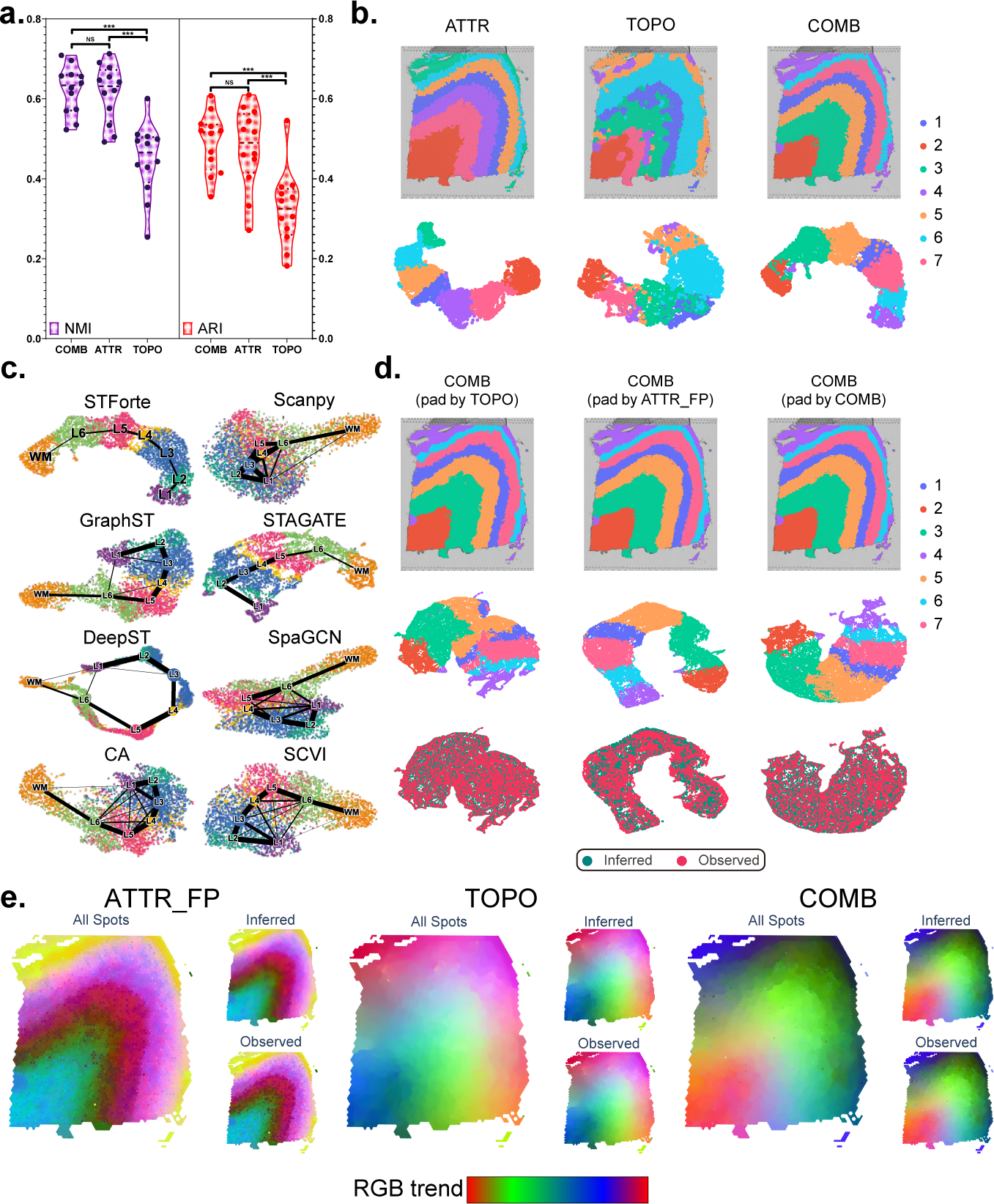
**a,** Violin plot of spatial region identification performance across the 12 slices of the dataset using different STForte encodings, quantified by the ARI and NMI metrics. The dashed lines represent the quartiles and median across 12 points. **b,** Identified spatial regions (top) and UMAP visualizations (bottom) obtained from different STForte encodings. **c,** Trajectories based on different latent embedding approaches incorporated with PAGA. **d,** Spatial regions (top), UMAP visualizations based on region annotations (middle), and spot instances (bottom) of different STForte encodings under padding scenario. **e,** RGB analysis depicting the identical spatial patterns and fluency of latent spaces from different STForte encodings within slice No. 151673.

**Extended Data Fig. 6.**
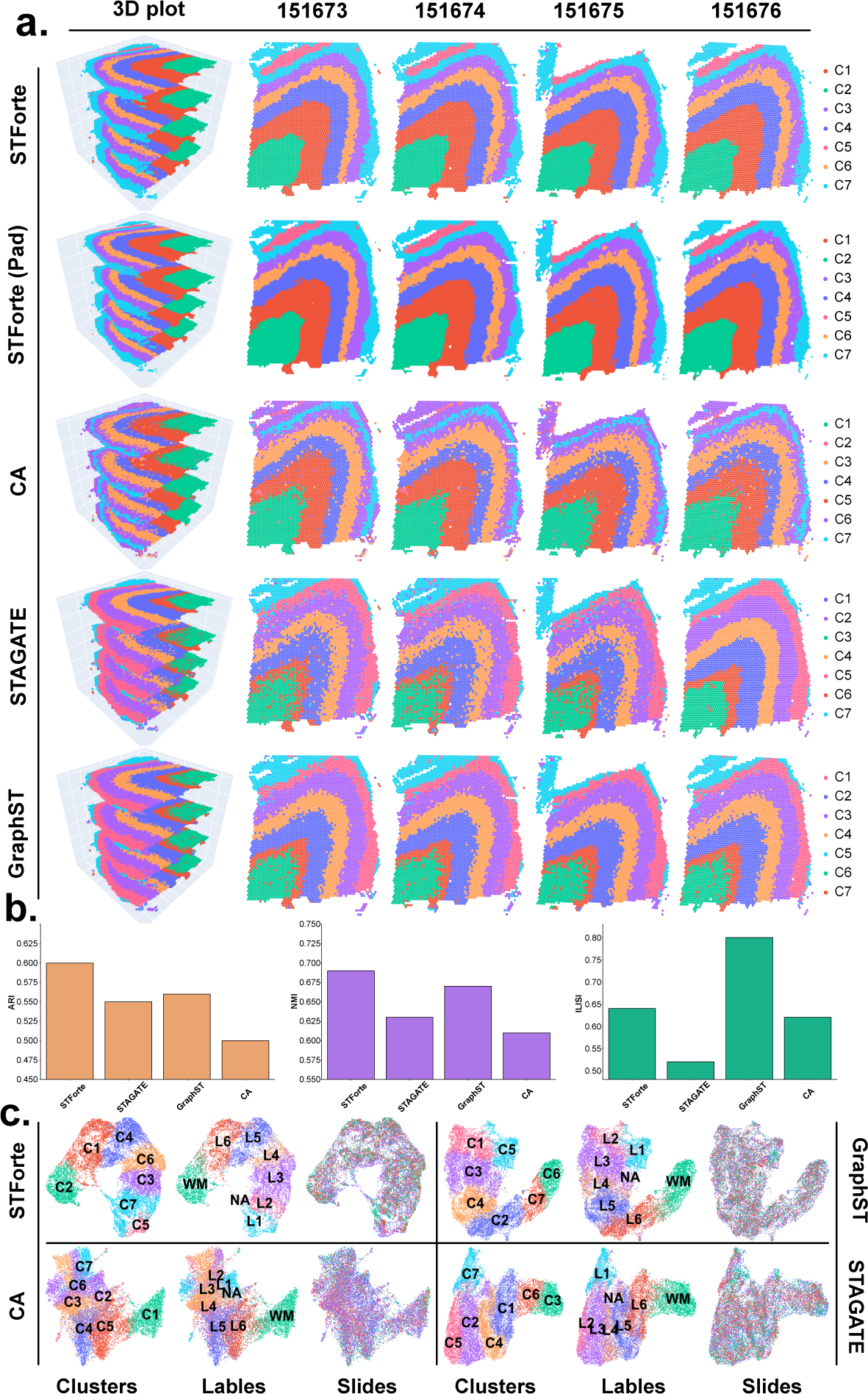
**a,** Spatial region identification results for the DLPFC dataset (No. 151673-151676) using different methods in multi-slice scenarios. **b,** Comparative performance of different methods in multi-slice analysis, including averaged ARI, averaged NMI, and iLISI. **c,** UMAP visualizations of multi-slice data obtained by different methods.

**Extended Data Fig. 7.**
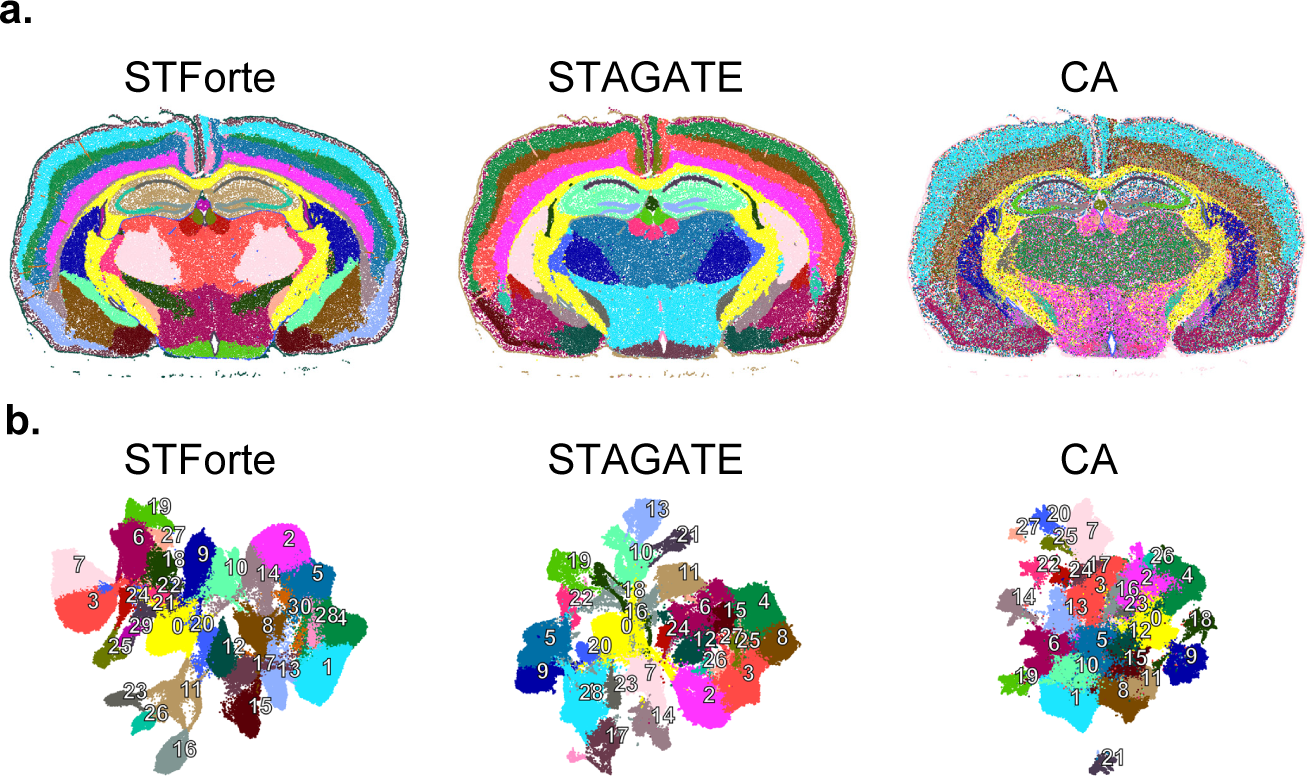
**a,** Spatial regions and **b,** UMAP visualization identified by different methods for 10x Xenium mouse coronal brain dataset.

**Extended Data Fig. 8.**
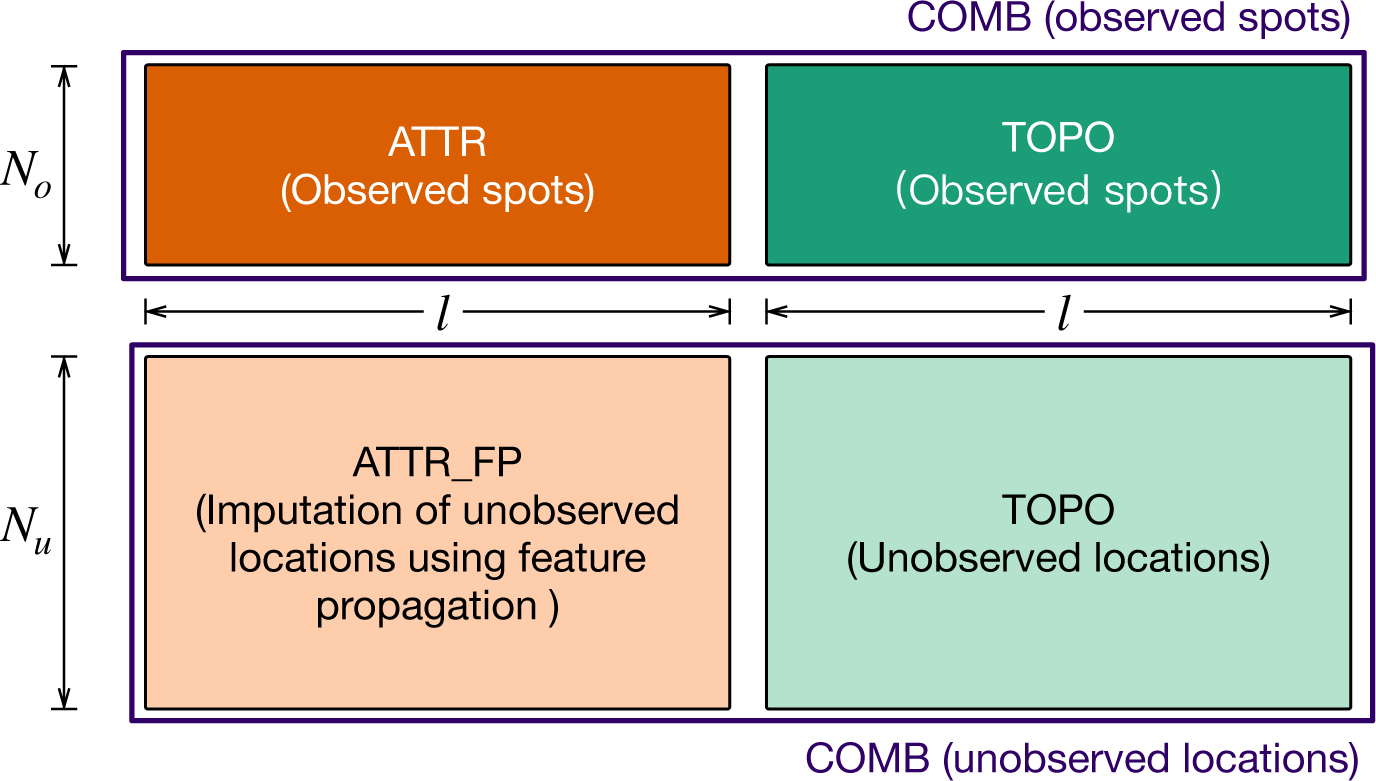
Schematic description of the encodings in the STForte model. The trained model generates attribute encoding (ATTR) and topology encoding (TOPO). ATTR contains only the patterns of observed spots, whereas TOPO additionally includes imputed information for unobserved locations. Furthermore, Feature propagation is performed to impute the ATTR encoding of unobserved locations. By concatenating ATTR and TOPO encodings, we obtain combined encodings (COMB), which provide more homogenized spatial information for downstream analysis compared to ATTR. TOPO demonstrates superior consistency between observed and unobserved locations, which facilitates its application for the property propagation of unobserved encodings. *N_o_*: data size of observed spots; *N_o_*: data size of unobserved locations; *l*: dimension of each latent encoding.

