## Supplementary Materials for "Enhanced spatially resolved transcriptomics analysis by matching between expression profiles and spatial topology"

### Supplementary Figures

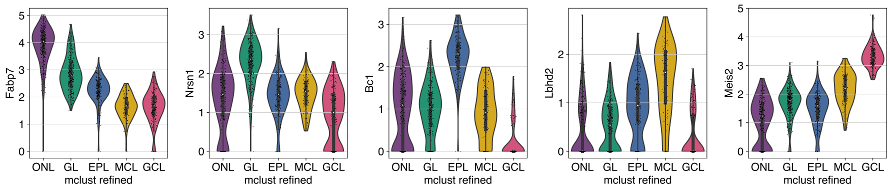


**Supplementary Fig. S1** A detailed violin plot shows the expression levels of different layer-specific marker genes across various layers.

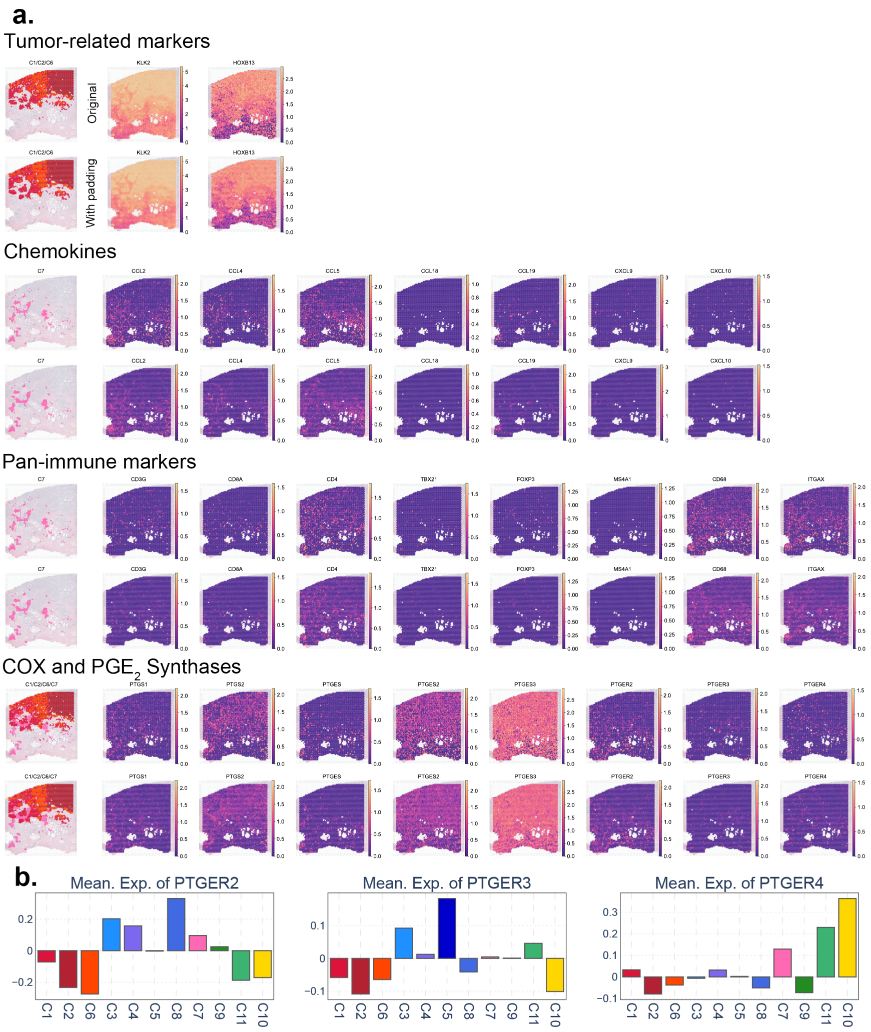


**Supplementary Fig. S2 a,** Investigated the spatial expression levels of cancer and immune-related genes, including results after STForte padding operation. **b,** Mean expression levels of different $PGE_2$ receptors.

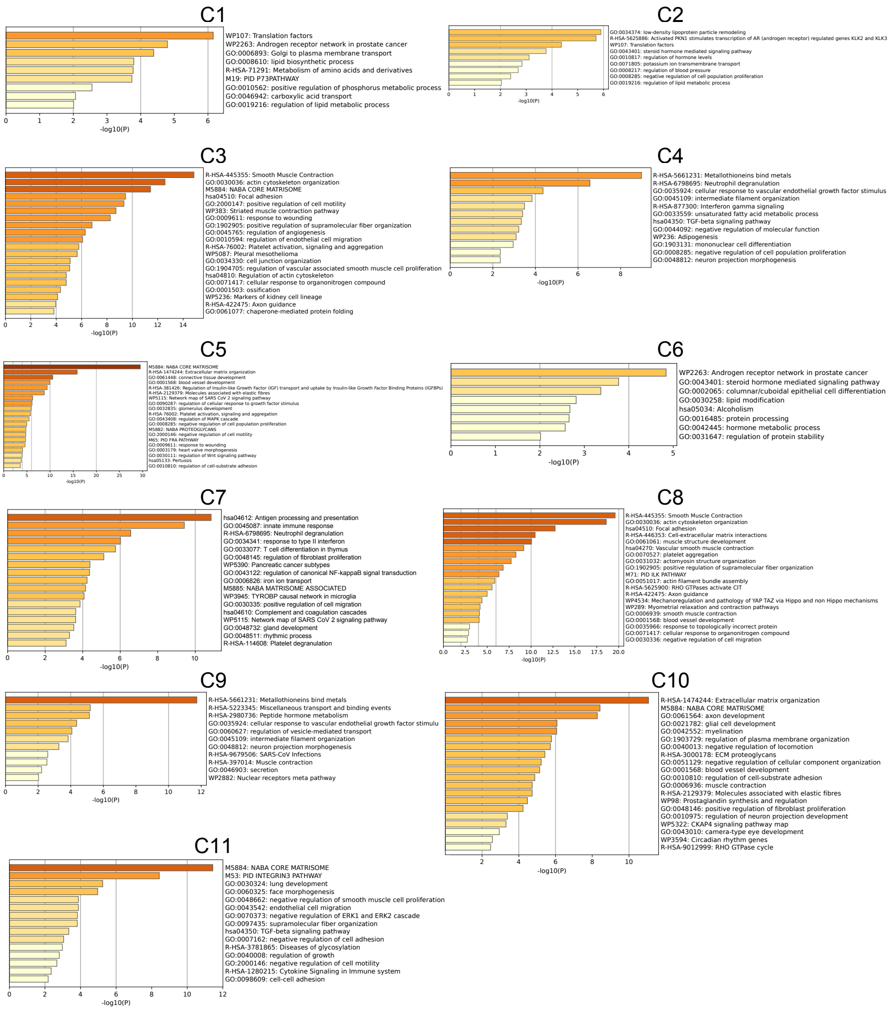


**Supplementary Fig. S3** Functional pathway analysis based on top differentially expressed genes specific to different spatial regions identified by STForte.

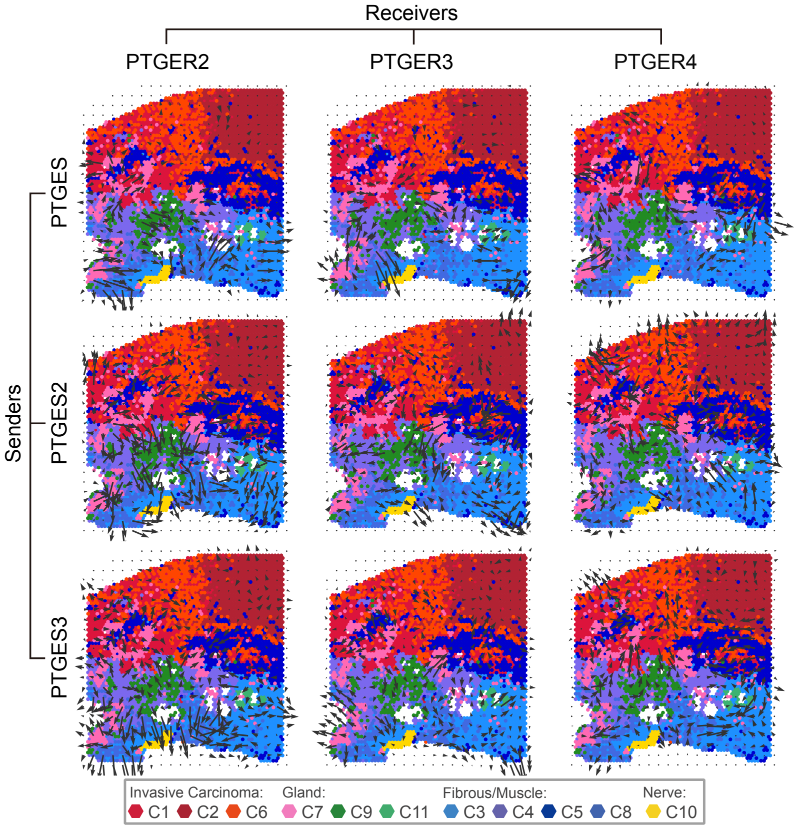


**Supplementary Fig. S4** The spatial interaction of relevant genes in the $PGE_2$ pathway, obtained by analyzing the results of spatial region identification using STForte in combination with COMMOT.

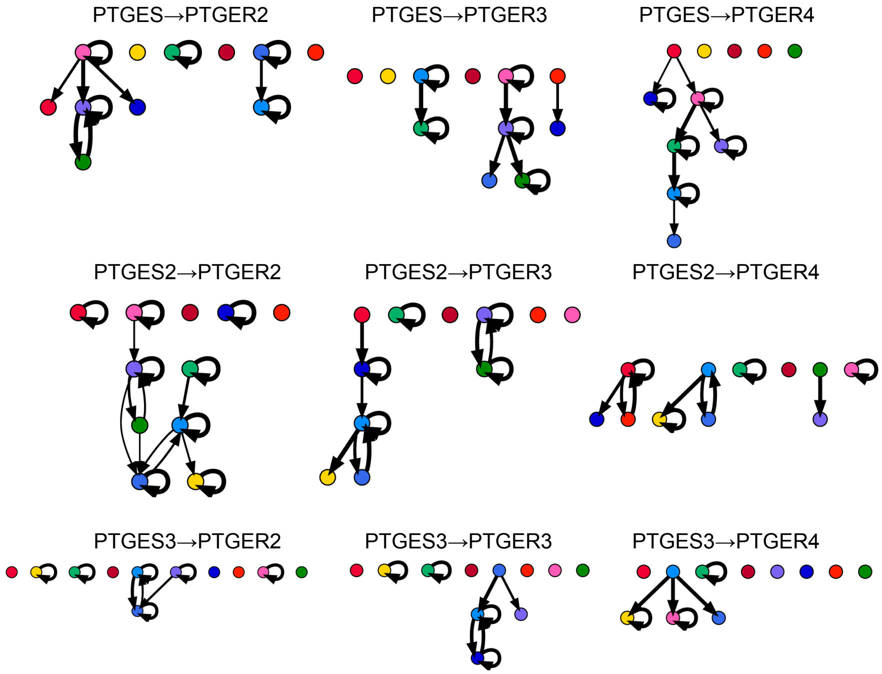


**Supplementary Fig. S5** The interaction among spatial regions of relevant genes in the $PGE_2$ pathway, obtained by analyzing the results of spatial region identification using STForte in combination with COMMOT.

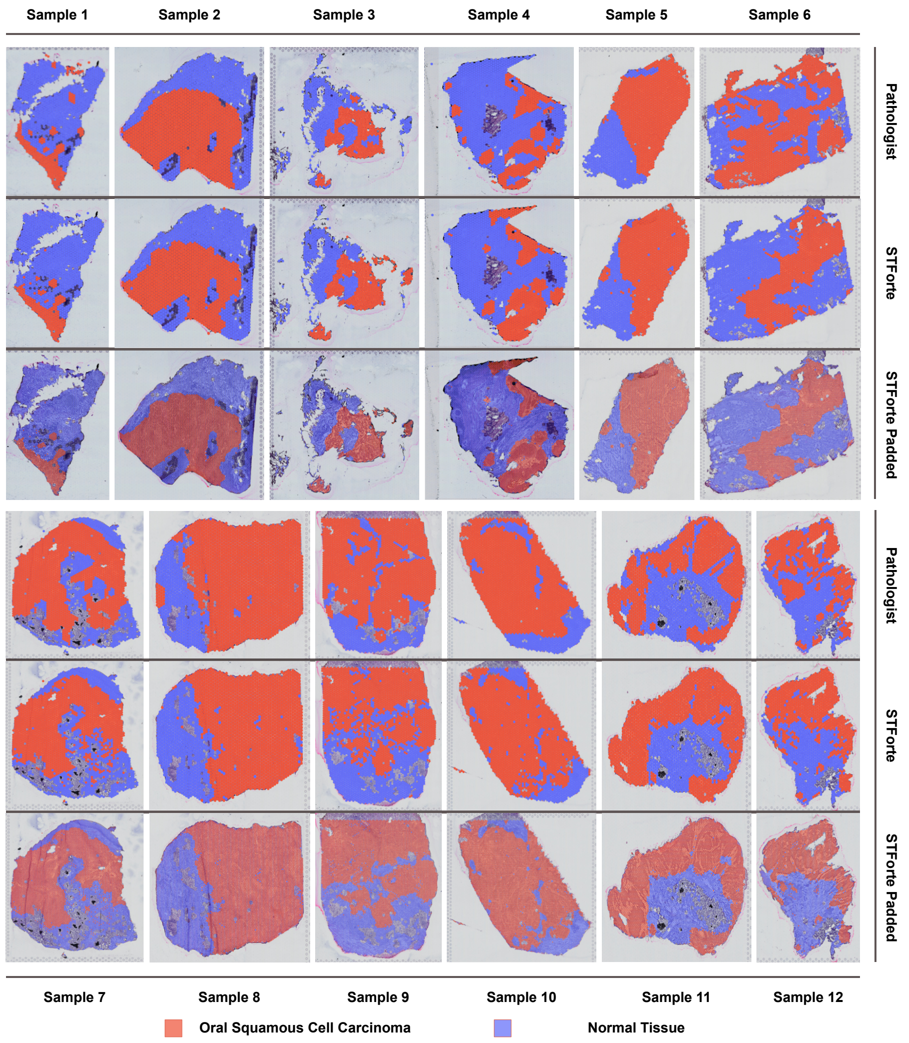


**Supplementary Fig. S6** Cancer region identification results of STForte for all 12 OSCC sections with padding. The SCC region was annotated through the Louvain algorithm with a resolution of 0.4 followed by manual cluster integration.

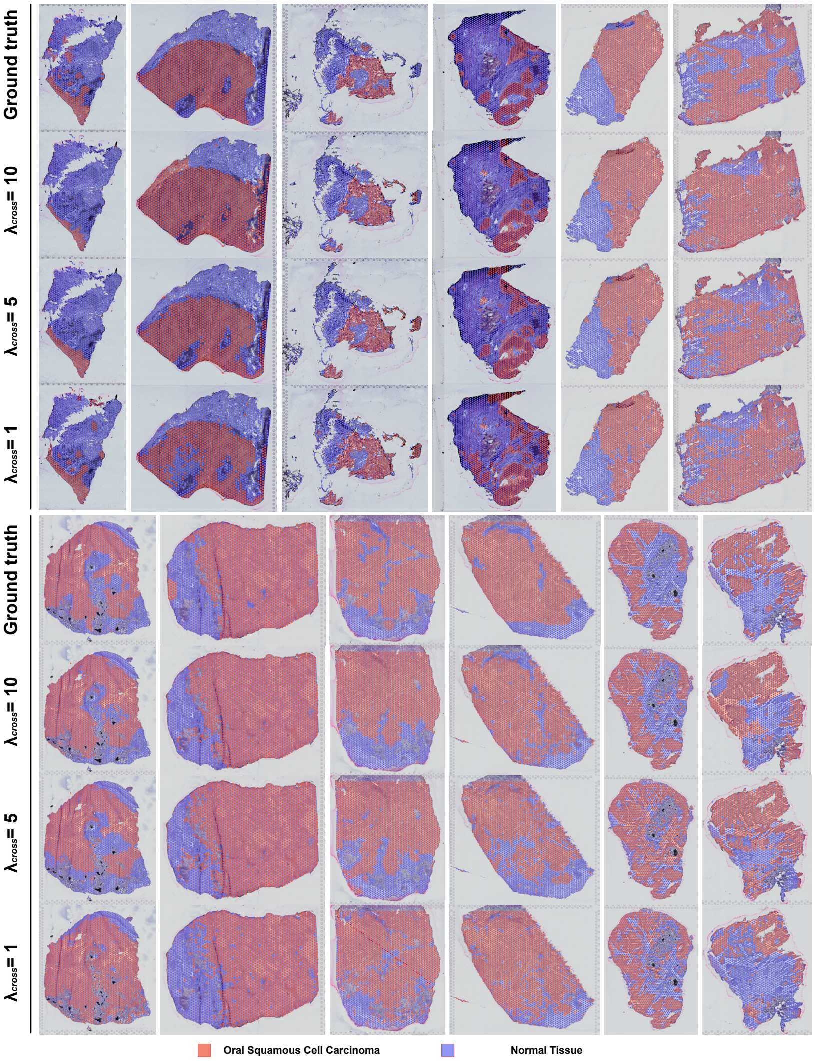


**Supplementary Fig. S7** Cancer region identification results of STForte under different settings of $\lambda_{cross}$.

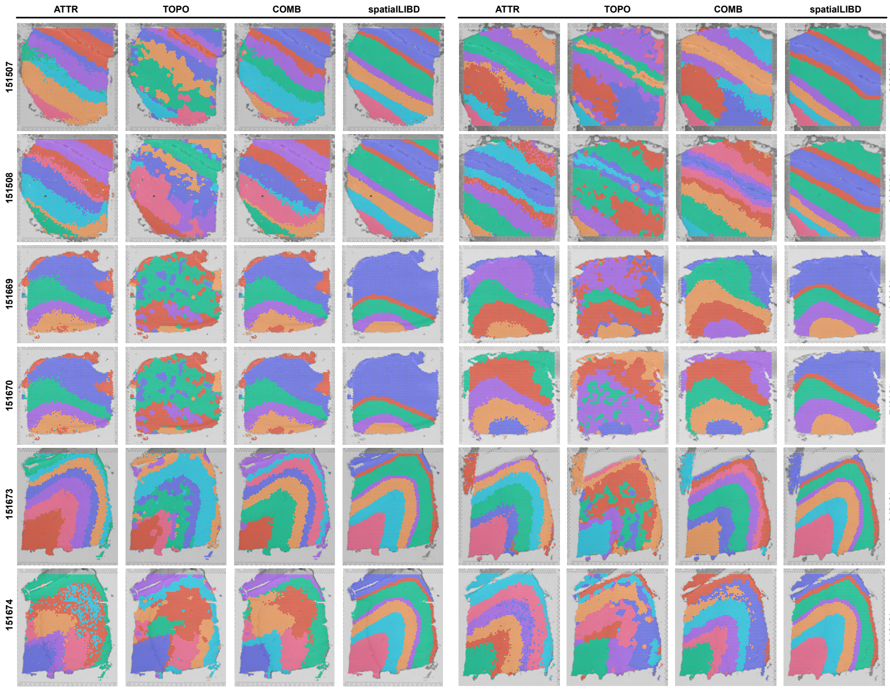


**Supplementary Fig. S8** Clustering results of all 12 sections of DLPFC.

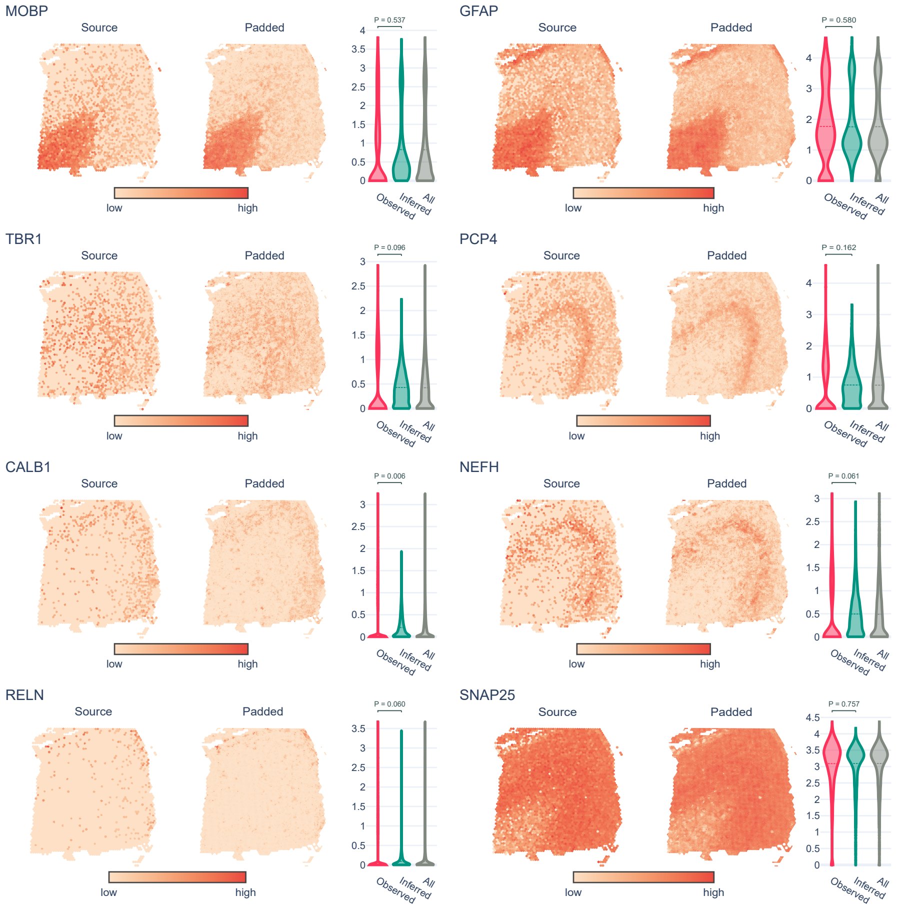


**Supplementary Fig. S9** Spatial gene expression levels of investigated layer-specific marker genes in the originally observed spots (source) and under the padding scenario. Violin plots depict the expression levels in observed spots, unobserved (inferred) spots, or all spots. P-values were obtained through t-tests.

### Supplementary Tables

| | ONL | GL | EPL | MCL | GCL | Excluded |
|---|---|---|---|---|---|---|
| **Fabp7 (ONL)** | 1.00E + 00 | 6.48E − 48 | 6.32E − 60 | 2.04E − 44 | 9.62E − 65 | 2.75E − 98 |
| **Nrsn1 (GL)** | 3.47E − 41 | 1.00E + 00 | 2.71E − 39 | 1.43E − 28 | 1.89E − 52 | 1.74E − 71 |
| **Bc1 (EPL)** | 2.84E − 45 | 2.06E − 52 | 1.00E + 00 | 1.17E − 33 | 7.86E − 52 | 4.52E − 70 |
| **Lbhd2 (MCL)** | 7.16E − 23 | 3.12E − 22 | 2.85E − 09 | 1.00E + 00 | 8.01E − 24 | 6.36E − 25 |
| **Meis2 (GCL)** | 8.51E − 67 | 2.18E − 64 | 1.66E − 53 | 5.82E − 35 | 1.00E + 00 | 1.66E − 86 |

**Supplementary Table S1** One-sided Wilcoxon rank-sum test p-values of 10x Visium mouse olfactory bulb data. Rank-sum tests are conducted for the layer-specific marker gene of corresponding layer in interest compared with other clusters or the entire set excluded the layer in interest. The results show significant divergence for the expression level of marker genes (p-values < 1E-20).

| | ONL(5) | GL(3) | EPL(0) | MCL/IPL(4) | GCL1(1) | GCL2(2) | RMS(6) |
|---|---|---|---|---|---|---|---|
| **Fabp7** | 8.59E − 166 | 6.64E − 24 | ns | ns | ns | ns | ns |
| **Nrsn1** | ns | 1.47E − 34 | 8.68E − 12 | ns | ns | ns | ns |
| **Bc1** | ns | ns | 6.54E − 05 | 5.73E − 04 | ns | ns | ns |
| **Lbhd2** | ns | ns | ns | 8.87E − 08 | ns | ns | ns |
| **Pcp4** | ns | ns | ns | 1.73E − 13 | 9.48E − 280 | 4.46E − 110 | ns |
| **Gad1** | ns | ns | ns | 4.57E − 41 | 4.36E − 172 | 1.63E − 13 | ns |
| **Meis2** | ns | ns | ns | ns | 3.54E − 90 | 2.42E − 63 | 2.07E − 15 |
| **Mbp** | ns | ns | ns | ns | 7.76E − 11 | 2.35E − 08 | 1.42E − 37 |

**Supplementary Table S2** One-sided Wilcoxon rank-sum test p-values of Stereo-seq mouse olfactory bulb data. Rank-sum tests are conducted for the layer-specific marker gene under a specific layer compared with other layers (i.e., the entire set excluded the layer in interest). P-value $>$ 1E-3 will be considered as non-significant (ns).

| | ARI | | | NMI |
|---|---|---|---|---|
| | signed-rank | ranksum | signed-rank | ranksum |
| **COMB-ATTR** | 0.6221 | 0.9540 | 0.4697 | 0.8174 |
| **COMB-TOPO** | 0.0005 | 0.007 | 0.0005 | 0.0001 |
| **ATTR-TOPO** | 0.0049 | 0.0039 | 0.0005 | 0.0003 |

**Supplementary Table S3** P-values of Wilcoxon signed-rank tests and Wilcoxon rank-sum tests for the clustering performance metrics based on different STForte encodings. The investigation was conducted across 12 slices in the 10x Visium DLPFC dataset.

| Method | ARI | | NMI | |
|---|---|---|---|---|
| | signed-rank | ranksum | signed-rank | ranksum |
| **STAGATE** | 0.4697 | 0.4189 | 0.0522 | 0.1482 |
| **GraphST** | 0.5186 | 0.6861 | 0.0522 | 0.3263 |
| **BayesSpace** | 0.0001 | 0.0647 | 0.0161 | 0.2040 |
| **DeenST** | 0.0427 | 0.0833 | 0.3013 | 0.6422 |
| **SpaGCN** | 0.92 | 0.024 | 0.001 | 0.0003 |
| **CA** | 0.0342 | 0.0243 | 0.0001 | 0.0003 |
| **scVI** | 0.0005 | 0.008 | 0.0005 | 0.0001 |
| **scanpy** | 0.005 | 0.0001 | 0.0005 | $< 1e-6$ |

**Supplementary Table S4** P-values of Wilcoxon signed-rank tests and Wilcoxon rank-sum tests for the clustering performance metrics based on other methods versus STForte. The investigation was conducted across 12 slices in the 10x Visium DLPFC dataset.

| Cluster | Domain | Name |
|---|---|---|
| 1 | Cerebral cortex | L2/3 |
| 2 | Cerebral cortex | L6 |
| 4 | Cerebral cortex | L4 |
| 5 | Cerebral cortex | L5 |
| 8 | Cerebral cortex | Cortical subplate |
| 13 | Cerebral cortex | Piriform |
| 14 | Cerebral cortex | L6b |
| 15 | Cerebral cortex | Cortical amygdalar area (CAA) |
| 17 | Cerebral cortex | L1 |
| 28 | Cerebral cortex | |
| 30 | Cerebral cortex | |
| 9 | Cerebral nuclei | Striatum dorsal region |
| 10 | Cerebral nuclei | Striatum-like amygdalar nuclei |
| 0 | Fiber tracts | Fiber tracts |
| 11 | Hippocampal | Hippocampal stratum |
| 16 | Hippocampal | DG-sg ( Dentate gyrus, granule cell layer) |
| 23 | Hippocampal | CA1sp (Field CA1, pyramidal layer) |
| 26 | Hippocampal | CA2sp/CA3sp (Field CA2/3, pyramidal layer) |
| 6 | Hypothalamus | |
| 18 | Hypothalamus | Zona incerta |
| 19 | Hypothalamus | |
| 27 | Hypothalamus | [Subthalamic nucleus] |
| 12 | Meninges | |
| 20 | Meninges | |
| 21 | Meninges | |
| 3 | Thalamus | Polymodal association cortex related |
| 7 | Thalamus | Sensory-motor cortex related |
| 22 | Thalamus | Reticular nucleus of the thalamus |
| 24 | Thalamus | Epithalamus (Lateral habenula) |
| 25 | Thalamus | Epithalamus (Medial habenula) |
| 29 | Ventricular | |

**Supplementary Table S5** Region annotations of 10x Xenium mouse coronal brain dataset according to the spatial and anatomical characteristics derived from STForte spatial identification results.

| | L1 | L2/3 | L4 | L5 | L6 | L6b | C28 |
|---|---|---|---|---|---|---|---|
| **Nrep** | 1.00 | $< 1e-6$ | $< 1e-6$ | $< 1e-6$ | 1.00 | 1.00 | $< 1e-6$ |
| **Ccn2** | 1.00 | 1.00 | 1.00 | 0.24 | $< 1e-6$ | $< 1e-6$ | 1.00 |
| **Rprm** | 1.00 | 1.00 | 1.00 | $< 1e-6$ | $< 1e-6$ | 0.96 | 1.00 |
| **Fezf2** | 1.00 | 1.00 | 1.00 | $< 1e-6$ | $< 1e-6$ | $< 1e-6$ | 0.06 |
| **Rorb** | $< 1e-6$ | 1.00 | $< 1e-6$ | 1.00 | 1.00 | 1.00 | 1.00 |
| **Rasgrf2** | 1.00 | $< 1e-6$ | $< 1e-6$ | 1.00 | 1.00 | 1.00 | 1.00 |
| **Gfap** | $< 1e-6$ | 1.00 | 1.00 | 1.00 | $< 1e-6$ | $< 1e-6$ | 1.00 |

**Supplementary Table S6** One-sided Wilcoxon rank-sum test p-values of isocortex domain within 10x Xenium mouse coronal brain dataset. Rank-sum tests are conducted for the layer-specific marker gene under a specific layer compared with other layers (i.e., the entire set excluded the layer in interest).

| | **Hipp. Stratum** | **CA1sp** | **CA2sp/CA3sp** | **Dg-sg** |
|---|---|---|---|---|
| **Prox1** | 1.000 | 1.000 | 1.000 | $< 1e-6$ |
| **Neurod6** | 1.000 | $< 1e-6$ | $< 1e-6$ | 1.000 |
| **Wfs1** | 0.002 | $< 1e-6$ | 1.000 | 1.000 |
| **Cpne4** | 1.000 | 1.000 | $< 1e-6$ | $< 1e-6$ |

**Supplementary Table S7** One-sided Wilcoxon rank-sum test p-values of hippocampal domain within 10x Xenium mouse coronal brain dataset. Rank-sum tests are conducted for the marker gene under a specific region compared with other layers (i.e., the entire set excluded the region in interest).

### Supplementary Notes

### 1 Graph Construction and Padding Strategy

#### 1.1 Neighborhood Graph

STForte offers two graph construction approaches according to the context of different ST technologies. For technologies that generate regular or approximately regular spot lattices, such as Visium or Ståhl *et al.*, neighborhood graphs are preferred in their original resolution. In this case, the distance parameter $d$ corresponds to the distance between adjacent spot centers in the regular lattices.

In situations where the distance parameter is not provided or the data exhibits approximately regular spot lattices, STForte can estimate the spot distance by utilizing an expected number of neighbors $n$ and an error bound $\epsilon$. The estimation process involves randomly sampling a spot from the dataset and assessing the differences in distances between the nearest $n$ spots and the sampled spot. If all of these differences fall within the specified $\epsilon$, the maximum value of these $n$ distances is recorded. Conversely, if any of the differences exceed $\epsilon$, a failure trial is noted. This iterative process continues until a predetermined number of iterations is reached. If the majority of trials yield bounded differences, the highest recorded distance is considered the estimated spot distance.

With a defined or inferred value for $d$, the neighborhood graph is subsequently defined, wherein the neighbor set of a given spot $i$ consists of all spots located within a Euclidean circle of radius $d$ centered at the location of spot $i$, which is:

$$\mathcal{N}(i) = \{j; d(i,j) \leq d \ \forall j = 1, \ldots, N\}, \tag{1}$$

where $d(i,j)$ is the Euclidean distance between the locations of spot $i$ and $j$.

#### 1.2 KNN Graph

STForte applied KNN graphs for ST data with irregular lattice, which is a commonly encountered scenario in single-cell resolution datasets. In order to construct a KNN graph, a fixed value for $K$ which represents the number of neighbors must be specified. For each individual cell, Euclidean distance is measured between its location and the locations of all other cells within the dataset, and the $K$ closest cells are selected to form the neighborhood set of that particular cell. Mathematically speaking, the neighbor set of cell $i$ is defined as

$$\mathcal{N}(i) = \{j; d_{(k)}(i,j) \ \forall j = 1, \ldots, N \ and \ k \leq K\}, \tag{2}$$

where the $d_{(k)}(i,j)$ denotes the $k^{th}$ ordered distance.

#### 1.3 Padding Strategy

The capability of STForte to infer the expression profile of unobserved locations has inspired the exploration of super-resolution using the platform through manual generation of pseudo spots. The concept is based on the observation that in Visium, there is a $45\mu m$ gap between spots, approximately equivalent to the diameter of a spot. Thus, the insertion of pseudo spots between each neighboring spot pair represents a potential method for achieving higher resolution. This approach involves the insertions of pseudo spots at the midpoint of each edge of the neighborhood graph constructed following the process outlined in the Section 1.1 and then infer the expression profiles of those pseudo spots using STForte.

#### 2 Pathfinder discovery network

In order to encode the topology information, STForte utilizes a Pathfinder Discovery Network (PDN) [1] for the topology encoder. Specifically, denote $\mathbf{H} \in \mathbb{R}^{n \times d_{\text{in}}}$ as the input and $\mathbf{H}' \in \mathbb{R}^{n \times d_{\text{out}}}$ as the output of a PDN layer and $\tilde{\mathbf{A}} \in \mathbb{R}^{n \times n}$ is the weighted adjacency matrix, where $n$, $d_{\text{in}}$ and $d_{\text{out}}$ are the node size (i.e, the number of nodes), input dimension and output dimension, respectively. The message passing operation can be calculated as follows:

$$\mathbf{H}' = \hat{\sigma}(\mathbf{D}_{\hat{\mathbf{G}}}^{-\frac{1}{2}} \hat{\mathbf{G}} \mathbf{D}_{\hat{\mathbf{G}}}^{-\frac{1}{2}} \mathbf{H}\mathbf{W} + \mathbf{b}) \tag{3}$$

$$\hat{\mathbf{G}}_i = g_\Omega(\tilde{\mathbf{A}}_i) \tag{4}$$

where $g_\Omega$ is an multi-layer perceptron (MLP) operator. $\mathbf{D}_{\hat{\mathbf{G}}}$ is the diagonal degree matrix of $\hat{\mathbf{G}}$. $\mathbf{W}$ is a learnable weight matrix and $\mathbf{b}$ is the bias vector. $\hat{\sigma}$ is an activation function.

While STForte accommodates the input of spatial distances among spots (cells) as a weighted adjacency matrix, it is important to note that in its default configuration, only the neighbor information between spots (cells) are utilized as input for the adjacency matrix (i.e., $\mathbf{A} \in \{0, 1\}^{n \times n}$). Under these circumstances, the PDN is functionally equivalent to a Graph Convolutional Network (GCN) [2], with calculation as follows:

$$\mathbf{H}' = \hat{\sigma}(\mathbf{D}^{-\frac{1}{2}} \hat{\mathbf{A}} \mathbf{D}^{-\frac{1}{2}} \mathbf{H}\mathbf{W} + \mathbf{b}) \tag{5}$$

where $\hat{\mathbf{A}} = \mathbf{A} + \mathbf{I}$ is the adjacency matrix with inserted self-loops. $\mathbf{D}$ is the diagonal degree matrix.

Moreover, to introduce nonlinearity and prevent overfitting, the rectified linear unit (ReLU) [3] activation function and dropout regularization [4] technique are incorporated into the graph neural network architecture.

### 3 Feature propagation

STForte also adopts a feature propagation (FP) [5] method to impute the unseen expression patterns from unobserved locations. FP intends to minimize the Dirichlet energy to achieve diffusion-based feature reconstruction through an iterative optimization. Specifically, $\mathbf{x} \in \mathbb{R}^N$ is a feature vector and $\mathbf{A} \in \{0,1\}^{N\times N}$ for a graph with N denotes the number of nodes in total. Following with the statements of STForte, we can assume that the $\mathbf{x}$ is composed by $\mathbf{x}_o \in \mathbb{R}^{N_o}$ for observed spots and $\mathbf{x}_u \in \mathbb{R}^{N_u}$ for unobserved spots. Therefore, the ordering of nodes are arranged as follows:

$$\mathbf{x} = \begin{bmatrix} \mathbf{x}_o \\ \mathbf{x}_u \end{bmatrix}, \mathbf{A} = \begin{bmatrix} \mathbf{A}_{oo} & \mathbf{A}_{ou} \\ \mathbf{A}_{uo} & \mathbf{A}_{uu} \end{bmatrix} \tag{6}$$

where $\mathbf{A}$ is as a partitioned matrix with different blocks are considered as the sub-adjacent matrix for observed-observed nodes ($\mathbf{A}_{oo}$), observed-unobserved nodes ($\mathbf{A}_{ou}$ and $\mathbf{A}_{uo}$ with $\mathbf{A}_{ou}^\top = \mathbf{A}_{uo}$) and unobserved-unobserved nodes ($\mathbf{A}_{uu}$), respectively. To this end, denote the normalized adjacent matrix as $\mathbf{A}' = \mathbf{D}^{-\frac{1}{2}}\mathbf{A}\mathbf{D}^{-\frac{1}{2}}$ with $\mathbf{D}$ being the degree matrix of $\mathbf{A}$, we can iteratively update $\mathbf{x}$ as follows:

$$\mathbf{x}^{(k+1)} = \begin{bmatrix} \mathbf{I} & 0 \\ \mathbf{A}'_{uo} & \mathbf{A}'_{uu} \end{bmatrix} \mathbf{x}^{(k)} \tag{7}$$

where $\mathbf{A}'_{uo}$ is the sub-adjacent matrix for the connection from unobserved spots to observed spots and $\mathbf{A}'_{uu}$ represents the sub-adjacent matrix from the unobserved spots.

The FP method was implemented in two distinct parts throughout our experiments:

(i) Within the main infrastructure of STForte, FP was employed to perform initial interpolation of expression profiles at unobserved locations to learn the topology encoding.

(ii) In our comparative analysis, FP was applied to attribute encoding to generate corresponding interpolations for unobserved spots, thereby enabling evaluation and visualization of the representational capacity and imputation capabilities of different encodings.

### 4 Evaluation Metrics

This section discusses the formula and implementation of the metrics used in this study.

#### 4.1 Clustering metrics

This study utilized four metrics to quantify the clustering performance including Adjusted Rand Index (ARI), Normalized Mutual Information (NMI), Silhouette Coefficient (SC), Calinski-Harabaz Score (C-H), and Local Inver Simpson's Index (LISI). The first two metrics were used to demonstrate the clustering performance when the true cluster labels were known. The following two metrics were considered to quantify the performance of different methods in real exploratory data analysis. The last metric LISI was used to quantify the performance of multi-slices clustering results. The first four metrics were calculated using the function in the metric module of package Sci-kit Learn and the LISI was implemented via package scib through the interface provided by Scanpy.

##### 4.1.1 Adjusted Rand Index (ARI)

ARI is a statistical measure that quantifies the similarities between two clustering labels of the same dataset. ARI picks a value between -1 and 1, where 1 indicates perfect agreement and -1 implies the counterproductive result. The calculation of ARI extends the Rand Index (RI) by adjusting for the expected agreement that would occur by chance, which could be formalized as

$$ARI = \frac{RI - E[RI]}{Max(RI) - E[RI]} \tag{8}$$

where RI computes the similarities between two clustering labels by considering all pairs of samples and counting disagreement pairs, which is

$$RI = \frac{N_{agree}}{N_{total}} \tag{9}$$

Where $N_{agree}$ denotes the number of agreed assignments and $N_{total}$ indicates the total number of pairs between two clusters.

##### 4.1.2 Normalized Mutual Information (NMI)

NMI is another statistical measurement that quantifies the same goal as ARI. It serves as a normalized version of mutual information score (MI) by dividing the generalized mean of entropies of the two cluster assignments, which is defined as follows,

$$NMI(U, V) = \frac{2MI(U, V)}{H(U) + H(V)} \tag{10}$$

where $U$ and $V$ denote the two cluster assignments. $H(\dot{})$ calculates the entropy, and $MI(U,V)$ represents the mutual information between cluster assignments. As a result, NMI ranges from 0 to 1, where a higher value indicates a better agreement between two cluster assignments.

###### 4.1.3 Silhouette Coefficient (SC)

SC [6] measures the separation among clusters which takes a value from -1 to 1. A higher value of SC indicates the clusters are more separated from each other which implies a more reasonable clustering result. Given the clustering assignment, SC first computes the intra-cluster distance denoted by $d_{itra}$ and the minimum of inter-cluster distance denoted by $d_{inter}$. Then SIL is defined as

$$SC = \frac{d_{inter} - d_{intra}}{max(d_{inter}, d_{intra})} \tag{11}$$

###### 4.1.4 Calinski-Harabaz Score (C-H)

The C-H, also known as the Variance Ratio Criterion, is a statistical measure of the separation of clusters [7]. It is particularly useful when comparing clustering performance when the number of clusters of clustering labels is different. Essentially, the C-H compares the ratio of the distance between the cluster centers to the average distance of points within a cluster. The formula of the C-H is

$$C - H = \frac{tr(S_b)}{tr(S_w)} \times \frac{n-k}{k-1} \tag{12}$$

where $S_b$ is the between-cluster scatter matrix, $S_w$ is the within-cluster scatter matrix, $n$ is the total number of samples, and $k$ is the number of clusters. A higher value of C-H indicates that the clusters are well-defined and well-separated from each other.

###### 4.1.5 Integration Local Inverse Simpson's Index (iLISI)

Local Inverse Simpson's Index (LISI) [8] is an extension of Simpson's Index (SI) which could be calculated as

$$LISI = \frac{1}{\sum_{i=0}^{K} p_i^2} \tag{13}$$

where $p_i$ is the proportion that the cluster label $i$ is in the local neighborhood, and $K$ is the total number of some categorical effects. LISI accesses whether cells are well-mixed across some categorical variables, like batch, technologies, and donor. Traditional LISI takes a value from 0 to number of batches and the iLISI used in this study scales the result to a number from 0 to 1 but does no affect the trend of LISI. This study utilized iLISI to compare the batch removal performances of different methods. A higher value of iLISI indicates a better mixture of batches which means better batch removal results.

#### 4.2 Other Metrics

STForte as a representation learning model with spatial enhancement ability, this study used multiple metrics to quantify its downstream analysis abilities, including Mean Squared Error (MSE), and Accuracy Score (ACC). This study used MSE to numerically demonstrate the performance of recovering gene expression of unobserved spots using different methods. ACC was calculated to assess whether these methods could precisely recognize the spots that should have non-zero expression value by recovering their expression level $\geq 0.5$ These two metrics were calculated through the metrics module of Scikit-Learn.

### 5 Compared Methods

#### 5.1 STAGATE

STAGATE [9] is a computational model designed for spatial transcriptomics data analysis, integrating gene expression with spatial information through a graph attention auto-encoder (GAE) framework to identify spatial domains and denoise data. Different from the traditional GAE framework, STAGATE replaces the outer-product decoder by a transposed GNN module whose parameters are copied from the encoder to reconstruct the gene expression. Experimentally, this study followed the official tutorial (https://stagate.readthedocs.io/en/latest/) to train STAGATE over 10xVisium and multi-slices 10x Visium data. The results demonstrated using Stereo-seq and Xenium data imitated the procedure for analyzing Slide-seq data with refined radius cutoff to ensure each spot gets approximately 10 neighborhoods on average.

#### 5.2 GraphST

GraphST [10] proposed a self-supervised graph contrastive learning framework for exploiting spatial information together with gene expression profiles. It is one of the current benchmarks of spatial domain identification. The clustering results of 10x Visium data produced in this study follow the procedures presented in the official tutorial of GraphST (https://deepst-tutorials.readthedocs.io/en/latest/). The multi-slide results were done by applying Paste2 to align the spatial locations first before getting into the official procedure.

#### 5.3 BayesSpace

BayesSpace [11] is a fully Bayesian statistical approach that ingeniously utilizes neighborhood structures in spatial transcriptomic data, allowing for accurate clustering, differential expression analysis, and enhancement of spatial resolution. In this study, the DLPFC clustering result was produced exactly following the same settings in its original tutorial (http://www.ezstatconsulting.com/BayesSpace/articles/maynard_DLPFC.html). The spatial resolution enhancement results were calculated using the same procedure and parameter settings in the demonstration of SCC (http://www.ezstatconsulting.com/BayesSpace/articles/ji_SCC.html ).

#### 5.4 DeepST

DeepST [12] is also a graph-based representation learning method that incorporates the GAE framework with an additional expression decoder. DeepST also provides a more fine-grained procedure for expression preprocessing and constructing the spatial graph. The clustering result of the DLPFC of DeepST was constructed under the instructions shown on its GitHub pages (https:

//github.com/JiangBioLab/DeepST) with slight modifications due to the different versions of package dependencies.

#### 5.5 SpaGCN

SpaGCN [13] integrates spatial information, gene expression, and histological image via graph convolutions and produces comprehensive embeddings for SRT data. Like STForte, SpaGCN uses the top PCA components as model input instead of dealing with raw counts directly. In the comparison part of this study, the result of SpaGCN was produced following its tutorial (https://github.com/jianhuupenn/SpaGCN/blob/master/tutorial/tutorial.md) but without the consideration of histological image for fairness (All other methods do not need histological information for embeddings).

## 5.6 CA

The Correspondence Analysis (CA) [14] is not a popular method in traditional single-cell and SRT data analysis. However, due to our analysis experience, this method is simple but usually efficient and effective. The CA is a matrix factorization method that serves as an alternative to PCA on count data. Different from PCA which approximates the Euclidean distance for differences in embeddings, CA performs the decomposition over the overall chi-squared statistic matrix. As a result, CA could theoretically avoid the unsatisfied continuous Gaussian assumption of PCA. The original article of CA provides source codes in R and this study translated the canonical form CA into Python codes and embedded it in the STGraph module.

#### 5.7 scVI

scVI [15] is a deep-learning-based non-spatial embedding method that is popular for dealing with single-cell data. It adopts a variational auto-encoder framework with considerations of library size and batch effects. This study followed the guidelines of the package scvi-tools (https://docs.scvi-tools.org/en/stable/tutorials/index.html) to generate the clustering result by treating the SRT data as traditional single-cell data.
